# Goal-directed recruitment of Pavlovian biases through selective visual attention

**DOI:** 10.1101/2022.04.05.487113

**Authors:** Johannes Algermissen, Hanneke E.M. den Ouden

## Abstract

Prospective outcomes bias behavior in a “Pavlovian” manner: Reward prospect invigorates action, while punishment prospect suppresses it. Theories have posited Pavlovian biases as global action “priors” in unfamiliar or uncontrollable environments. However, this account fails to explain the strength of these biases—causing frequent action slips—even in well-known environments. We propose that Pavlovian control is additionally useful if flexibly recruited by instrumental control. Specifically, instrumental action plans might shape selective attention to reward/ punishment information and thus the input to Pavlovian control. In two eye-tracking samples (N = 35/ 64), we observed that Go/ NoGo action plans influenced when and for how long participants attended to reward/ punishment information, which in turn biased their responses in a Pavlovian manner. Participants with stronger attentional effects showed higher performance. Thus, humans appear to align Pavlovian control with their instrumental action plans, extending its role beyond action defaults to a powerful tool ensuring robust action execution.

## Public significance statement

This study suggests that Pavlovian biases, a fast-and-frugal decision strategy that may trigger suboptimal choices in certain contexts, are not a permanent, immutable force upon behavior. Instead, they are flexibly recruited depending on the action a person is planning: Given the plan to make/ withhold actions, people preferably attend to reward-/ punishment-related information, which in turn trigger Pavlovian biases that facilitate the implementation of these plans. Stronger “outsourcing” of action implementation to such an attentional recruitment of Pavlovian biases leads to higher performance. These findings highlight how Pavlovian biases are more flexible than previously thought and how strong biases can be of advantage.

Goal-directed recruitment of Pavlovian biases through selective visual attention The valence of potential outcomes biases action selection: The prospect of rewards invigorates action (“Go”), while the prospect of punishment suppresses it (“NoGo”). These so-called motivational, or “Pavlovian”, biases have first been observed in animal studies in which the presence of a reward-associated cue invigorated cue-unrelated behaviors (Estes, 1943, 1948; LoLordo et al., 1974; Lovibond, 1983; Schwartz, 1976). While at first interpreted as seemingly irrational, recent theorizing has suggested that these biases in fact constitute a decision-making strategy that is particularly “fast-and-frugal” (Boureau et al., 2015; Dayan et al., 2006). Past theorizing has assumed that, while inflexible, these biases are fast, computationally cheap, and likely attuned to global environmental statistics (Dayan et al., 2006). They can thus act as sensible “default” response strategies in situations in which instrumental, goal-directed control fails to deliver rewards beyond chance levels, such as novel or uncontrollable environments (Daw et al., 2005; Dorfman & Gershman, 2019; O’Doherty et al., 2017). These accounts assume that Pavlovian and instrumental control co-exist, largely segregated from another, and merely compete at the behavioral output level. In case of conflict, the former has to be actively suppressed—a requirement humans only imperfectly master (Breland & Breland, 1961; Cavanagh et al., 2013; Hershberger, 1986; Swart et al., 2018).

Several fields within psychology, including research on decision-making, motor control, and attention, have recognized that, in order to solve a given problem, an agent can use different strategies. To pick a situationally appropriate strategy, it does not only matter whether or how well the strategy solves the problem (e.g., how many rewards it returns), but also what the invested costs are (e.g., how long it takes, how many mental resources it takes) (Bettman et al., 1990; Boureau et al., 2015). Hence, seemingly suboptimal or “irrational” behavior can turn out to be rational when seen in the light of costs or resource constraints—a term called “bounded rationality” (Simon, 1957) or more recently “resource rationality” (Griffiths et al., 2015; Lieder & Griffiths, 2020). Strategies that return higher-quality solutions in some situations might become unfeasible in other situations due to resource constraints, calling for simpler, less costly strategies. This viewpoint has the potential to not only explain the choice of seemingly inferior options violating the axioms of rational choice (Palminteri et al., 2015; Tversky, 1969), but also motor errors (Du et al., 2022; Hardwick et al., 2019; McDougle et al., 2016; Wolpert & Landy, 2012) and seemingly nonstrategic, imprecise, or inefficient (“lazy”) visual search (Araujo et al., 2001; Ballard et al., 1995; Draschkow et al., 2021; Horowitz & Wolfe, 1998; Steinman et al., 2003; Wolfe et al., 2000). In all these psychological domains, humans (and other animals) have seemingly multiple independent decision-making systems at their disposal.

One class of particularly simple decision strategies are so-called “heuristics” or “biases”, fast-and-frugal decision strategies which are rather inflexible, but perform well in a restricted set of situations (Gigerenzer & Gaissmaier, 2011; Hutchinson & Gigerenzer, 2005). These heuristics have likely been acquired as adaptations to specific environmental challenges through both biological and cultural evolution (Haselton et al., 2009; Todd & Brighton, 2016). The fact that many heuristics are present even in animals (Fawcett et al., 2014) speaks for their evolutionary ancientness and possibly genetic hardwiring. However, the ”meta”-question arises how to determine which heuristic to use in a given situation (Lieder & Griffiths, 2017; Marewski & Link, 2014; Rieskamp & Otto, 2006). Apparently, humans and other animals frequently misapply heuristics (in such cases typically called “biases”) (Beck et al., 2012; Fawcett et al., 2014; Rahnev & Denison, 2018), raising the question why these biases are so seemingly strong and hard to suppress.

A relevant question in many fields of psychology is whether distinct strategies operate in isolation, conflict with each other, or even work in synergy. Specifically, more sophisticated strategies might “outsource” certain sub-routines to simpler strategies, yielding a “division of labor”. Such a synergy is frequently assumed to evolve over time, with initial acquisition through more “explicit” rule-driven strategies, which are later outsourced to more “implicit”, incremental, habit-like strategies, a division prominent in goal-directed vs. habitual decision-making (Balleine & Dickinson, 1998; Daw et al., 2005), response preparation (Du et al., 2022; Hardwick et al., 2019), explicit vs. implicit motor skill learning (Mazzoni & Krakauer, 2006; McDougle et al., 2016; McDougle & Taylor, 2019), and goal-directed vs. history-guided attention (Anderson, 2016; Theeuwes, 2018). Beyond such a sequential labor division in which one system hands over control to another, there are even examples of systems that are active simultaneously, where one system trains the other, e.g. in reward revaluation (Gershman et al., 2014; Robinson & Berridge, 2013), credit assignment (Moran et al., 2019), and memory replay (Mattar & Daw, 2018). Crucially, such a simultaneous collaboration requires both systems to be permanently active. In this paper, we propose that also instrumental and Pavlovian control can work in such a synergy.

In contrast to previous literature that has assumed a parallel, strictly segregated arrangement of instrumental and Pavlovian control, we suggest that the instrumental system can adaptively recruit and steer the Pavlovian system by selecting its input via visual attention. Humans are not just passively exposed to reward and punishment cues that drive these biases. Instead, they can actively seek out or ignore these cues and thereby modulate their influence via selective visual attention (“active sensing”) (Friston et al., 2010; Gottlieb & Oudeyer, 2018; Yang et al., 2016). In a world full of distractions, where actions unfold over time and are prone to interference, instrumental control could harness the power of cue-driven, “automatic” behavioral tendencies by directing visual attention to cues that activate them and then automatically trigger the intended action. In this scenario, it might be warranted to keep the Pavlovian system permanently “online”, accepting a few infrequent errors at the benefit of overall more robust action implementation. This view contrasts with earlier assumptions that Pavlovian biases are merely “defaults” to fall back to in novel or uncontrollable environments. Instead, keeping Pavlovian control constantly online during instrumental goal pursuit might be advantageous. However, previous task designs measuring Pavlovian biases do not match such scenarios in which agents actively seek out information that helps them achieve their goals. We developed a new paradigm that temporally separates action selection, attention to reward and punishment information, and action execution. We then tested whether humans seek out reward and punishment information—and allow Pavlovian biases to shape responding—in a way that is aligned with their action goals. Note that, in the following, we will use the term “goal-directed” to denote such a synchronization between action goals and information search—remaining tacit about whether the underlying cognitive process involves prospective planning or devaluation sensitivity, features typically taken as indicators of “goal-directedness” of behavior (Balleine & Dickinson, 1998).

Research in the past decade supports the notion that overt attention (eye gaze) towards positive aspects of choice options predicts their eventual selection (Cavanagh et al., 2014; Fiedler & Glöckner, 2012; Krajbich et al., 2010), while attention to negative aspects predicts their rejection (Armel et al., 2008; Pachur et al., 2018; Westbrook et al., 2020). In these studies, positive and negative information is required for making the correct choice. Theoretical perspectives have speculated that longer attention to an option facilitates memory retrieval of its features, which could accentuate its value (Shadlen & Shohamy, 2016; Weilbächer et al., 2021). However, attention to task-irrelevant positive or negative cues—which have no apparent relationship to the choice options and thus cannot serve as anchors for memory retrieval—might have similar effects. Indeed, in Pavlovian-to-Instrumental-Transfer (PIT) paradigms, incidental background cues associated with positive/ negative outcomes induce Go/ NoGo actions (Estes, 1943, 1948; Geurts et al., 2013a, 2013b; Huys et al., 2011; Rescorla & Soloman, 1967). Linking those PIT effects to the role of attention in value-based choice implies that directing attention to (task-irrelevant) reward or punishment cues should activate the Pavlovian system and, in this way, automatically invigorate or suppress choice.

Beyond effects of attention on action, there is also evidence that action plans themselves can direct attention (Heuer et al., 2020; Olivers & Roelfsema, 2020; van Ede, 2020). Task goals modulate which stimulus features we are sensitive to and distracted by (Eimer & Kiss, 2008; Folk et al., 1992; van der Stigchel & Hollingworth, 2018). “Active sensing” perspectives frame attention as a tool to actively interrogate the environment while implementing action plans (Cisek & Pastor-Bernier, 2014; Gottlieb & Oudeyer, 2018; Yang et al., 2016). The premotor theory of attention goes as far as proposing that the primary purpose of attention is to monitor target features relevant for preparing an action towards the target (Rizzolatti et al., 1987; Sheliga et al., 1997). Studies have indeed found perceptual sensitivity to be selectively sharpened for features relevant for an ongoing action, e.g. object location for reaching movements or object size and orientation for grasping movements (Bekkering & Neggers, 2002; Craighero et al., 1999; Fagioli et al., 2007). However, in the domain of value-based decision-making, similar evidence for task goals shaping attention is scarce. One relevant finding might be that humans tend to seek out a choice option one final time before selecting it (“last fixation” or “late onset” bias) even if they already know this option to be superior to other options (Hunt et al., 2016; Kaanders et al., 2021). In this case, attention appears to be guided by choice rather than vice versa, extending of the premotor theory of attention to value-based decision-making.

Taken together, there appear to be mechanisms synchronizing agents’ attention with their action plans, and there is tentative evidence for attention to reward and punishment information triggering automatic responses in the fashion of Pavlovian biases. Hence, it seems indeed possible that an instrumental system could “recruit” the Pavlovian system to “aid” the execution of action plans by strategically steering attention toward relevant information. We tested this idea in two samples (the second one a direct, pre-registered replication) using eye-tracking. For this purpose, we designed a novel Go/ NoGo learning task in which action planning and execution were separated by a phase in which participants could preview the positive or negative outcomes at stake. Notably, information about these outcomes was not informative for the selection of the correct action. We predicted that action plans would shape attention to reward and punishment stakes, i.e., that participants’ first fixation (not confounded by bottom-up saliency effects due to a gaze-contingent design) would be more often on the reward information when participants planned a Go (compared to a NoGo) action. Vice versa, we predicted an effect of attention duration to rewards vs. punishments on the final response, i.e., that longer attention to reward compared to punishment information would lead to more Go responses and speed up reaction times (Fig. 1A, B). Such a goal-directed recruitment of Pavlovian biases would extend their role beyond mere “default” strategies in novel environments towards a powerful aiding robust action execution.

**Figure 1.**
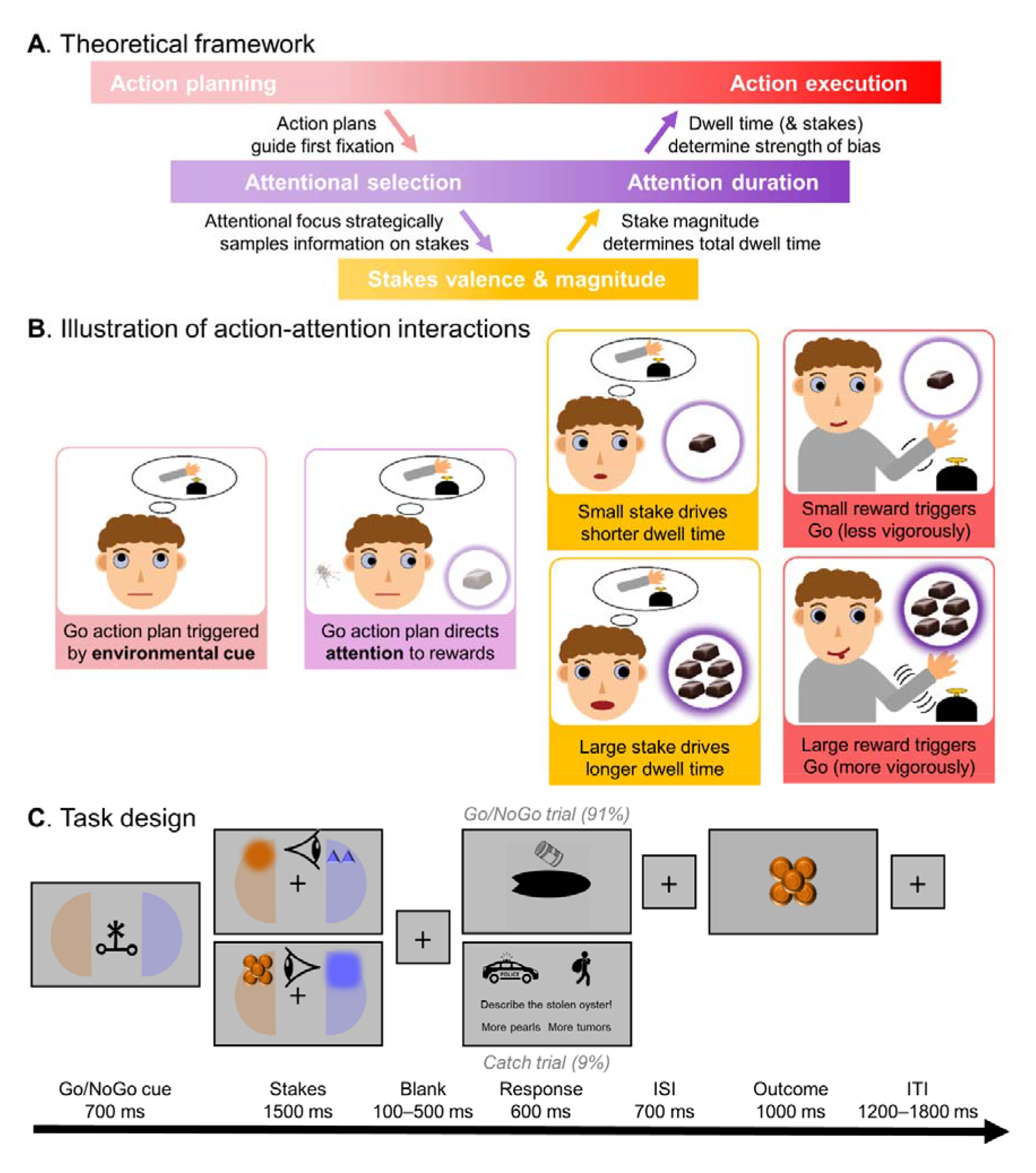
Theoretical framework and task design. **A.** Theoretical framework of the interaction between action and attention. An environmental cue elicits an action plan, which directs top-down attention (first fixation) towards information about potential reward/ punishment outcomes (stakes). The first fixation anchors attention and (partly) determines which stakes will receive more attention, which is additionally modulated by bottom-up signals such as the magnitude of the stakes. The relative attention on reward versus punishment stakes (dwell time) biases the final Go/ NoGo action in a Pavlovian manner. **B.** Cartoon illustration of the proposed interaction of action planning and attention. **C.** Task design. Participants learned Go/ NoGo responses to various cues (cover story: feed/ not feed various oyster types to maximize pearls and minimize toxic tumors). Cue presentation (instructing the correct action) and action execution are separated by a phase in which rewards (pearls, here orange) and punishments (toxic tumors, here blue) at stake for correct/ incorrect responses are presented in a gaze-contingent manner. Afterwards, the oyster (black oval) can be fed, and for Go responses, participants have to press the button on the side where it is “still open”. Outcomes are delivered in a probabilistic manner (75% feedback validity). On catch trials, participants have to indicate whether the oyster featured more pearls or tumors (cover story: The oyster is stolen by thieves and has to be retrieved back from the police, which requires identification).

## Methods

### Participants and Exclusion Criteria

In Sample 1, we recorded eye-tracking data from 35 participants (*M*_age_ = 23.7, *SD*_age_ = 4.1, range 18– 35, one outlier at age 58; 27 women, 8 men; 30 right-handed; 21 with the right eye as dominant eye). In Sample 2 (replication sample), we recorded data from 64 participants (*M*_age_ = 21.5, *SD*_age_ = 3.0, range 18–34; 50 women, 13 men, 1 other; 62 right-handed; 41 with the right eye as dominant eye). In this replication sample, the study design, hypotheses, and analysis plan were pre-registered (https://osf.io/nsy5x). The sample size for this sample was based on the effect size of the primary effect of interest in Sample 1, i.e., action requirements affecting first fixations (*z* = 2.89, Cohen’s *d* = 0.49), which yielded a required sample of N = 57 to detect such an effect with 95% power (two-sided one-sample t-test) (Murayama et al., 2022). We initially collected data from 57 participants, but given that seven participants did not perform significantly above chance level, we collected additional seven participants. Performance above 56% in 240 trials was significantly above chance (one-sided binomial test). Note that, in line with our pre-registration, all results in the main text are based on all participants (see Supplemental Material 1 for an overview of all results); results for only those participants that performed significantly above chance are reported in Supplemental Material 2 and led to identical conclusions.

Participants were recruited via the SONA Radboud Research Participation System of Radboud University. Exclusion criteria comprised glasses, color blindness, and prior treatment for neurological or psychiatric disorders. The study protocol was identical for both samples. Participants took part in a 1h session that comprised informed consent, eye-tracker calibration, a 10-minute practice phase including written instructions and practice trials, and finally the 30-minute eye-tracking experiment. Upon completion of the task, participants filled in a structured debriefing about their presumed hypothesis of the experiment, and any strategies they applied. None of the participants guessed the study hypotheses. Participants received a participation fee of €10 or 1h of course credit plus a performance dependent-bonus of €0–2 (Sample 1: *M* = €0.77, *SD* = €0.43, range €0.09–1.58; Sample 2: *M* = €0.91, *SD* = €0.47, range €0.10–1.67). Research was approved by the local ethics committee of the Faculty of Social Sciences at Radboud University (proposal no. ECSW-2018-171).

### Apparatus

Reporting follows recently suggested guidelines for eye-tracking studies (Fiedler et al., 2020). The experiment was performed in a dimly lit, sound-attenuated room, with participants’ head stabilized with a chin rest. The experimental task was coded in PsychoPy 2020.2.7 on Python 3.7.0, presented on a 24’’ BenQ XL2420Z screen of resolution (1920 x 1080 pixels resolution, refresh rate 144 Hz). Manual button presses were applied via a custom-made button box with two buttons (index and middle finger of the dominant hand). Participants’ dominant eye was tracked with an EyeLink 1000 tracker (SR Research, Mississauga, Ontario, Canada; sampling rate of 1,000 Hz; spatial resolution of 0.01° of visual angle, monocular recording), controlled via Pylink for Python 3.7.0. The eye-tracker was placed 20 cm in front of the screen, and participants’ chin rest 90 cm in front of the screen. Before the task, participants performed a 9-point calibration and validation procedure (software provided by SR Research). Calibration was repeated until an error < 1° was achieved for all points. The screen background grey tone (RGB 180, 180, 180) was constant across calibration and the experimental task.

### Task

Participants performed a Go/ NoGo learning task with delayed response execution, called the Oyster Farming Task (Fig. 1C). On each trial, participants cultivated an oyster that could either grow 1–5 pearls or 1–5 hazardous tumors. Pearls gained money while tumors cost money for disposal. To maximize the probability that oysters grew pearls, participants needed to learn which oysters to “feed” (Go) and which ones not to feed (“NoGo”). Crucially, participants could choose to reveal the reward (number of pearls) and punishment (number of tumors) at stake prior to action execution in a gaze-contingent design. Participants’ score of accumulated money was turned into a bonus of 0–2€ at the end of the task. Participants performed 264 trials split into three blocks of 88 trials (80 trials of the Go/ NoGo task, 8 catch trials), each with a new set of four oyster types. For detailed information on the instructions, see the original materials used in this study available in the data sharing collection under [All data and code will be made available upon manuscript acceptance].

Each trial started with one (of four) abstract *action cues* (letters from the Agathodaimon alphabet; size 5.2° x 5.2°) presented for 700 ms in the center of the screen, representing an oyster type. For each oyster type, there was an optimal action (feed or not feed) that participants needed to learn by trial-and-error. Feeding was only possible when the oysters “opened” later in the trial. The optimal action led to rewards (pearls) in 75% of (valid) trials, otherwise to punishments (tumors; on “invalid trials”). Vice versa, suboptimal actions led to punishments on valid trials, but to rewards on invalid trials. During action cue presentation, participants were informed about the sides (left vs. right) on which upcoming stakes information (rewards vs. punishments) would appear via faintly colored semi-circles in the respective colors (blue and orange, counter-balanced across participants).

Directly after action cue off-set, participants were cued with the exact locations of the stakes and given 1,500 ms to unveil the tumors and pearls at stake on the respective trial. Stakes were revealed in a gaze-contingent fashion: fuzzy circular color patches appeared on the semi-circles, which changed into the number of pearls/ tumors at stake when participants fixated them. This eliminated any bottom-up saliency effects (e.g., of stake magnitude) on peripheral vision that could affect participants’ first fixations. To prevent exact pre-programming of saccades, exact locations of stakes varied across trials. Stakes were located on an invisible circle with a radius of 5.2° visual angle around the screen center (i.e., distance of stakes from the center was kept constant), with a potential vertical displacement of −45 – +45 degrees from the horizontal midline. Vertical displacement was always identical for both pearls and tumors. Stakes were represented by circular areas of interest (AOI) of 150 pixels (2.7°), with a minimal distance between stakes (at maximal vertical displacement) of 514 pixels (9.4°) and a maximal distance (positioned on the horizontal midline) of 852 pixels (15.6°). Stakes were presented in orange (RGB 200, 100, 7) and blue (RGB 104, 104, 255) of equal luma. Stakes varied in magnitude (1–5 items; total display size 2.6° x 2.6°) and magnitude was balanced within action cues (i.e., each of the 20 possible combinations used once per cue, excluding the five possible combinations in which both magnitudes were identical). The mapping of pearls and tumors to the left/ right side varied across trials and was balanced within action cues (each side 10 times per cue) to control for possible participant-specific side biases in gaze.

Stakes offset was followed by a variable interval of 100–500 ms (uniform distribution in steps of 100 ms), after which a release cue (black oyster shape and a food can, 5.2° x 5.2°) appeared for 600 ms, indicating that the oyster was about to close and could be fed if necessary. The oyster remained open at either the left or right side, indicating the side where the oyster could be fed. If participants chose to feed the oyster, they had to press the respective button on the open side. The uncertainty about the response side (left/ right) at the time of the action cue, which was only resolved with the release cue, prevented pre-mature responding. In-time responses were confirmed by the food can (1.7° x 1.7°) tipping over to the respective side. 700 ms after release cue offset, the outcome (3.5° x 3.5°) was presented for 1,000 ms. Late responses during the release cue-outcome interval were recorded, but did not affect the outcome. Pressing the incorrect button (i.e., the oyster was open on the left/ right, but participants pressed the right/ left button) counted as incorrect (i.e., yielded tumors on valid trials) and was confirmed by the can tipping over in the respective direction. Participants received a number of either pearls or tumors, depending on the stakes, their response, and trial validity. Trials finished with a variable inter-trial interval between 1,200 and 1,800 ms (uniform distribution in steps of 100 ms).

On 8 out of 88 trials per block, participants performed a catch task which incentivized attention to the stakes: Instead of the release cue, participants had to report whether the reward or punishments stakes were of greater magnitude (Fig. 1D). These catch trials encouraged participants to monitor both stakes and process their magnitude.

### Data Preprocessing

#### Behavior

Catch trials were excluded from all analyses of responses and RTs. We further excluded trials with RTs below 200 ms (% trials with button presses per participant: Sample 1: *M* = 0.1, *SD* = 0.3, range 0–1.5; Sample 2: *M* = 0.2, *SD* = 0.3, range 0–1.1) because such fast responses could not be expected to incorporate processing of the cue. Likewise, we excluded trials RTs above 800 ms (% trials with button presses per participant: Sample 1: *M* = 0.9, *SD* = 1.6, range 0–6.8; Sample 2: *M* = 0.5, *SD* = 1.8, range 0–14.0). This deadline was 200 ms after release cue offset (i.e., closing of the response window) as we reasoned that any later responses could have been triggered by the release cue offset. Go responses with the incorrect hand were very rare (% trials with incorrect hand response per participant: Sample 1: *M* = 1.7, *SD* = 3.1, range 0–14.6; Sample 2: *M* = 1.3, *SD* = 2.4, range 0–13.3) and not significantly influenced by stakes or dwell times.

#### Eye-tracking preprocessing

Gaze data was processed in R with custom-code. Continuous data was epoched into trials of 1500 ms relative to stakes onset. Gaps of missing samples up to a duration of 75 ms (due to blinks or saccades) were interpolated using linear interpolation. Trials with more than 50% of missing samples were discarded altogether (% trials per participant: Sample 1: *M* = 4.5, *SD* = 8.0, range 0–34.1; Sample 2: *M* = 3.5, *SD* = 7.9, range 0–52.7). Gaze position was marked as being on the reward/ punishment stakes when gaze position was less than 150 pixels away from the center of the respective stakes image, which was also the criterion in our gaze-contingent design for rendering stakes visible. For each trial, we computed the first fixation on any stakes object (reward or punishment) as well as the total duration (in ms) with which rewards and punishments were fixated over the entire trial (“dwell time”). Absolute dwell times were converted into dwell time difference (reward time minus punishment time) and dwell time ratios (reward time divided by the sum of reward and punishment time) (Westbrook et al., 2020).

On some trials, participants only fixated one stake (% trials with at least one fixation per participant: Sample 1: *M* = 11.0, *SD* = 14.6, range 0–61.4; Sample 2: *M* = 10.0, *SD* = 14.4, range 0– 50.4), leading to ratios of 0 or 1. We thus deviated from our pre-registration and report results for dwell time difference (reward minus punishment dwell time) in the main text, which avoids such an accumulation of values at the edges; analyses of dwell time ratio are reported in Supplemental Material 1 and led to identical conclusions. Analyses using only the trials on which participants fixated both stakes led to largely identical conclusions.

### Data Analysis

#### General strategy

We tested hypotheses using mixed-effects linear regression (function lmer) and logistic regression (function glmer) as implemented in the package lme4 in R (Bates et al., 2015). We used generalized linear models with a binomial link function (i.e., logistic regression) for binary dependent variables such as responses (Go vs NoGo) and first fixation, and linear models for continuous variables such as RTs or dwell time. We used zero-sum coding for categorical independent variables. All continuous dependent and independent variables were standardized such that regression weights can be interpreted as standardized regression coefficients. All regression models contained a fixed intercept. We added all possible random intercepts, slopes, and correlations to achieve a maximal random effects structure (Barr et al., 2013). *P*-values were computed using likelihood ratio tests with the package afex (Singmann et al., 2018). We considered *p*-values smaller than α = 0.05 as statistically significant.

The main analyses were pre-registered for Sample 2 (replication sample; pre-registration available under https://osf.io/nsy5x). We deviated from our pre-registration by reporting results based on dwell time differences (reward minus punishment dwell time) instead of dwell time ratios (reward dwell time divided by reward plus punishment dwell time) in the main text. When participants fixated only one stake, the dwell time ratios were either 0 or 1, regardless of the absolute dwell time on each single fixated option, leading to a loss of information and an accumulation of values at the edges, yielding a distribution with three modes. In contrast, dwell time differences were approximately normally distributed and statistically more comparable to stake differences. Nonetheless, analyses of dwell time ratio and dwell time differences led to identical conclusions as reported in the Supplemental Material 1.

#### Baseline learning and Pavlovian biases

First, following previously established motivational Go-NoGo learning tasks (Guitart-Masip et al., 2011; Swart et al., 2017), we tested i) the degree to which participants learned the task, i.e., performed more Go responses to Go cues than NoGo cues, and ii) whether responses were influenced by the magnitude of the reward and punishment stakes, reflecting the presence of a Pavlovian bias. For this purpose, we fitted mixed-effects regressions with responses (Go/ NoGo) and (as secondary variable) reaction times as dependent variables and a) the required action (Go/ NoGo) as well as b) the difference in reward and punishment stake magnitude (ranging from −4 to +4) as independent variables. A significant effect of stake difference was followed up with post hoc analyses separating the effects of reward and punishment stake magnitudes, reported in Supplemental Material 3.

#### Analysis of gaze patterns

Our first key prediction was that action plans, elicited by the oyster cues, directed attention towards action-congruent stakes (reward stake for Go requirement, punishment stake for NoGo requirement). The crucial test of this prediction was whether the action requirement elicited by the cue affected the location of the first fixation (on the reward versus the punishment stake). This first fixation was not confounded by any bottom-up saliency effects since, in our gaze-contingent design, the magnitudes of the stakes was not visible yet. We used both required action (Go or NoGo) and the difference in the modeled Q-values for Go relative to NoGo responses as independent variables to predict the first fixation. These analyses also included catch trials since, during the stakes phase, participants were unaware of whether the trial would be a Go/ NoGo or catch trial. All eye-tracking analyses contained a regressor capturing any participant-specific side biases (overall preference to fixate the left or right).

#### Computational modelling of action values

We tested the impact of participants’ action intentions on their attention towards the reward and punishment stakes using two operationalizations: Firstly, we approximated participants’ intentions by the action required by the presented cue (oyster type). However, this operationalization assumes that participants (have learned and) know the required action. This assumption is violated i) at the beginning of blocks when participants cannot know the required action yet and still have to acquire it through trial-and-error, and ii) even more so in participants who fail to learn the correct response for (some of) the cues. Thus, secondly, as a more proximate measure of participants’ beliefs of what they should do, we fitted a simple Rescorla-Wagner Q-learning model to the Go/ NoGo response data of each participant. This model uses outcomes *r* (+1 for rewards, −1 for punishments; given that the exact outcome magnitude is irrelevant for learning) to update the action-value *Q* for the chosen action *a* towards cue *s*:

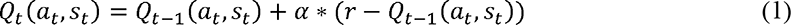

Action values were then translated to action probabilities using a Softmax choice rule:

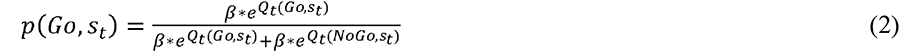

The model featured the free parameters α and β. The learning rate α determines the impact of prediction errors (i.e., higher α leads to stronger incorporation of recent outcomes and discounting of past outcomes). The inverse temperature β determines the stochasticity of choices (i.e., higher β leads to more deterministic choices in line with action values and lower β to more noisy, stochastic choices). Both parameters were estimated to each participants’ data using a grid search, with α in the range [0, 1] in steps of 0.01 (Sample 1: *M* = 0.07, *SD* = 0.08, range 0.01–0.35; Sample 2: *M* = 0.14, *SD* = 0.18, range 0.001–0.84) and β in the range of [1, 40] in steps of 0.1 (Sample 1: *M* = 8.27, *SD* = 8.21, range 1.0–32.7; Sample 2: *M* = 8.64, *SD* = 9.57, range 1.0–34.8). Starting values for Q_Go_ and Q_NoGo_ were set to 0. Using each participants’ best fitting parameters as well as their action and outcomes on each trial, we then simulated the action values for Go and NoGo responses on each trial using one-step-ahead predictions (Steingroever et al., 2014). We used the difference term Q_Go_ – Q_NoGo_ as more proximate measure of participants’ action intentions on each trial based on their past experience with each cue. On catch trials (on which participants did not make a Go/ NoGo response and did not receive feedback), Q-values were not updated, but were carried over from the last cue encounter. Similarly, Q-values were not updated on trials on which participants responded in the incorrect direction (i.e., pressed left when the oyster was open on the right or vice versa) since participants were instructed that such “directional” errors were always counted as incorrect. Feedback was thus not informative as to whether a Go or NoGo response would have been correct for this cue.

#### Analysis of effects of attention on responses and reaction times

Our second key prediction was that attention to the reward and punishment stakes would shape action execution. To test this prediction, we tested whether the dwell time difference (milliseconds spent on reward stakes minus milliseconds spent attending to punishment stakes) predicted responses (Go vs. NoGo) and response speed (RT, for Go responses only). These analyses excluded catch task trials (where responses did not relate to learning but to comparing stake magnitudes). All analyses involving responses or reaction times as dependent variable controlled for the required response as well as participant-specific side biases (overall preference to first fixate the left or right). Results did not change when controlling for the Q-value difference instead of the required response.

Note that, in our pre-registration for Sample 2, we mentioned the plan to fit reinforcement-learning drift diffusion models (RL-DDMs) to the combined choice and RT data. See Supplemental Material 3 for a discussion of why these models were unable to reproduce important qualitative patterns present in the empirical data, which was likely due to the tight response deadline and the NoGo response option.

#### Between-subjects correlations of accuracy

If humans synchronized their attention with their action plans such that Pavlovian biases would align with instrumental action requirements, one would expect this process to facilitate task performance and lead to higher accuracy. To test whether participants with stronger effects of attention on the final response indeed showed higher accuracy, we performed exploratory analyses by computing between-subjects correlations between overall task accuracy and i) the degree to which stake differences (reward minus punishment stake magnitude) affected responses as well as ii) the degree to which relative dwell time (reward minus punishment dwell time) affected responses. For this purpose, we refit the respective models on all participants, collapsing across both samples (total *N* = 99), and computed between-subjects correlations between participants’ percent correct responses and their respective regression coefficient (fixed + random effect extracted).

### Transparency and openness

We report how we determined our sample size, all data exclusions, all manipulations, and all measures in the study. All data, analysis code, and research materials are available under [All data and code will be made available upon manuscript acceptance]. The study design, hypotheses, and analysis plan for Sample 2 were pre-registered under https://osf.io/nsy5x. Data were analyzed using R, version 4.1.3 (R Core Team, 2022). Models were fitted with the package lme4, version 1.1.31 (Bates et al., 2015). Plots were generated with ggplot, version 3.4.2 (Wickham, 2016).

## Results

### Learning and Pavlovian biases

Overall, participants learned the Go/ NoGo task (% correct, Sample 1: *M* = 70.0, *SD* = 10.4, range 50.0–87.1; Sample 2: *M* = 73.4, *SD* = 13.2, range 36.3–91.7), performing significantly more Go responses to Go cues than NoGo cues (Sample 1: *b* = 1.08, 95% CI [0.88 1.27], χ^2^(1) = 53.19, *p* < .001; Sample 2: *b* = 1.27, 95% CI [1.09 1.44], χ^2^(1) = 89.19, *p* < .001; Fig. 2A). Participants also performed well in the catch trials (% correct: Sample 1: *M* = 85.8, *SD* = 10.1, range 56.5–100; Sample 2: *M* = 86.2, *SD* = 15.5, range 25.0–100; Fig. 2D). Five (seven) people in Sample 1 (2) did not perform significantly above chance (56% correct based on a 1-sided binomial test with 240 trials) in the Go/ NoGo task. In line with our pre-registration, we still included these subjects in all our analyses (for results without these participants, see Supplemental Material 2). To account for variability in learning, we estimated action (Q) values for each trial based on a Rescorla-Wagner learning model.

**Figure 2.**
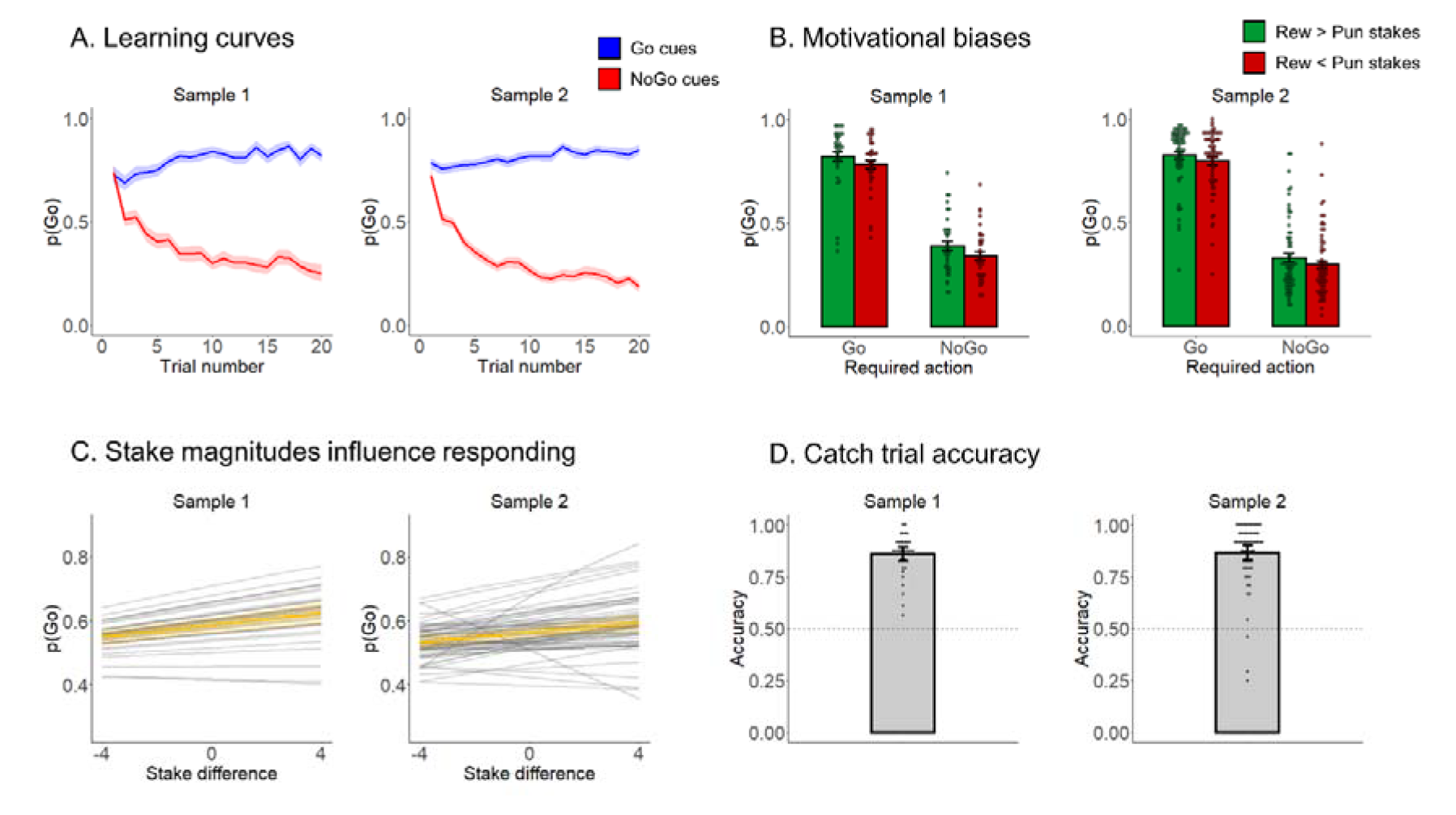
Task performance and Pavlovian biases. A. Performance in the Pavlovian Go/ NoGo task. Trial-by-trial proportion of Go actions (±SEM) for Go cues (blue lines) and NoGo cues (red lines). Shadows indicate standard errors for per-condition-per-participant means. Participants clearly learn whether to make Go actions or not (blue lines go up; red lines go down). B. Pavlovian biases. Participants perform more Go responses on trials where the reward stake was higher than the punishment stake (green bars) than vice versa (red bars). Individual data points reflect response proportion per participant. C. Stake magnitudes biased responding in a continuous fashion. A higher stake difference (i.e., a reward stake minus punishment stake) resulted in a higher proportion of Go responses. Faint grey lines represent regression lines per participant as predicted by the mixed-effects regression model; the bronze line represents the group-level regression line; bronze shading represent mean and 95% confidence intervals. Note the two strong outliers in Sample 2; excluding these outliers did not change conclusions. D. Performance in the catch trials. Individual data points reflect accuracy per participant.

Beyond outcome-based learning, responding was affected by the stake magnitudes in a way similar to previously observed Pavlovian biases. A more positive stake difference (reward minus punishment stake magnitude) increased the proportion of Go responses (Sample 1: *b* = 0.12, 95% CI [0.06 0.17], χ^2^(1) = 15.32, *p* < .001; Sample 2: *b* = 0.09, 95% CI [0.03 0.15], χ^2^(1) = 7.92, *p* = .005; Fig. 2B, C) and increased response speed (Sample 1: *b* = −0.04, 95% CI [-0.07 −0.01], χ^2^(1) = 7.32, *p* = .007; Sample 2: *b* = −0.03, 95% CI [-0.05 −0.004], χ^2^(1) = 6.31, *p* = .012). The effect of stakes differences did not become weaker over trials or blocks (see Supplemental Material 3). Separating these effects for the reward and punishment stakes showed that effects were driven by both valences: Higher (relative to lower) reward stake magnitude increased responding and speeded up responses, while higher (relative to lower) punishment stake magnitude decreased responding and slowed responses (see Supplemental Material 3).

In sum, we found evidence that participants learned the task and that the reward and punishment stake magnitudes biased responding in opposite directions, reflecting Pavlovian biases. For reaction times, we found larger reward stake magnitudes to speed up responding and larger punishment stake magnitudes to slow down responding, again in line with Pavlovian biases as reported in previous literature (Guitart-Masip et al., 2011; Swart et al., 2017).

### Action plans direct eye gaze

Next, we tested whether participants’ attention was synchronized to their action plans. Such a link would allow Pavlovian biases to be elicited specifically by reward/ punishment cues that trigger an action in line with participants’ intentions. As a measure of goal-directed attention, we used the first fixation on each trial (Konovalov & Krajbich, 2016), which was unaffected by any bottom-up saliency effects of the (yet to be uncovered) stakes in our gaze-contingent design. On trials that required a Go response, participants were significantly more likely to first fixate rewards than on trials that required a NoGo response (Sample 1: *b* = 0.11, 95% CI [0.04 0.19], χ^2^(1) = 13.92, *p* < .001; Sample 2: *b* = 0.09, 95% CI [0.03 0.15], χ^2^(1) = 7.88, *p* = .005; Fig. 3A).

**Figure 3.**
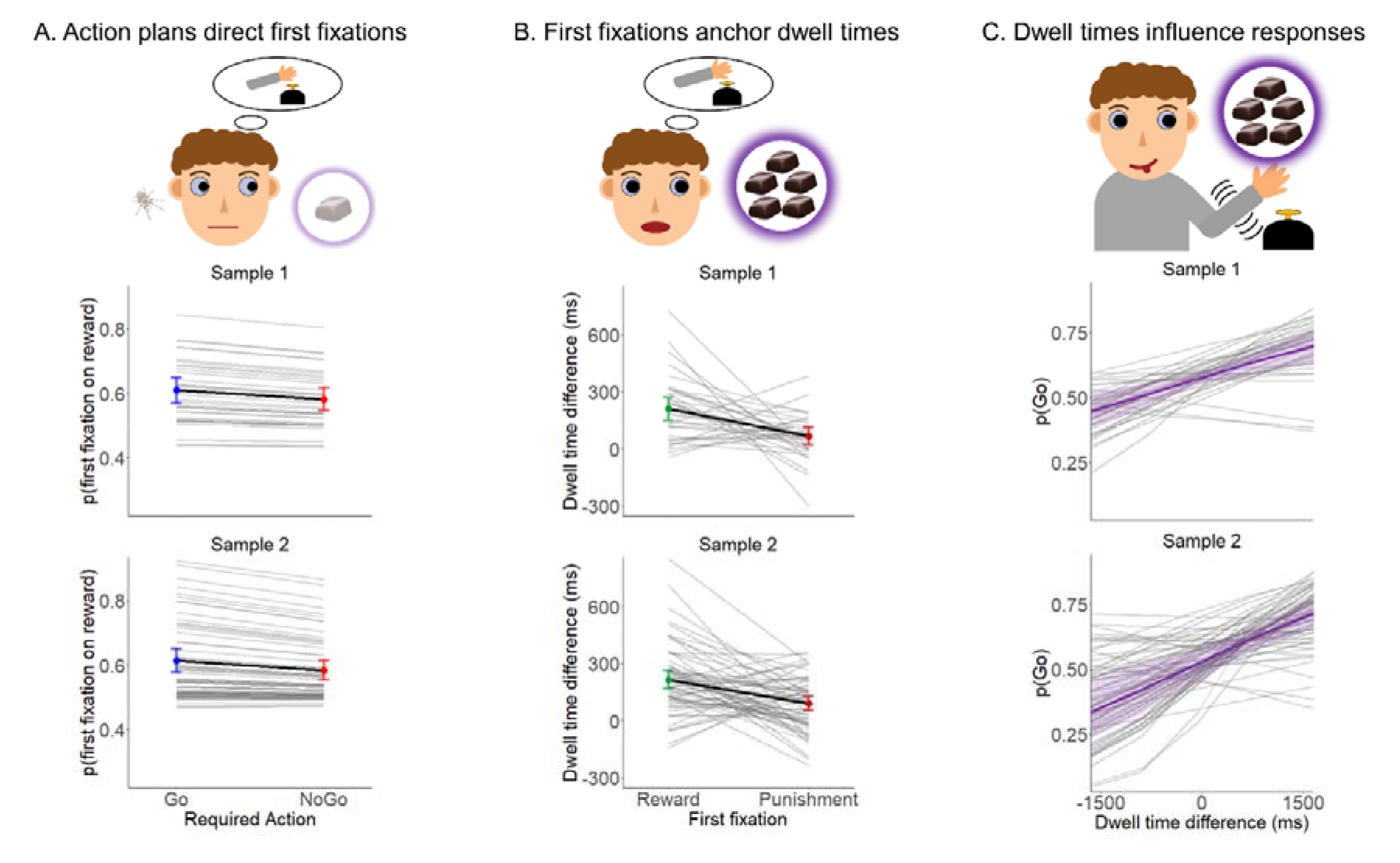
Mutual influences between action and attention. A. Action plans direct first fixations. When required to make a Go action, participants are more likely to first fixate reward information than when a NoGo action was required. B. First fixation anchors attention. Dwell times are longer on reward stakes compared to punishment stakes when the first fixation was already on reward stakes. Dwell times are additionally shaped by other factors such as the stake magnitudes. C. Dwell time differences affect final responses. Longer attention to reward compared to punishment stakes resulted in a higher proportion of Go responses. Grey lines = regression lines per participant as predicted by the mixed-effects regression model; Black line = group-level regression line; Shading = the 95% confidence interval.

This analysis used the required response as a predictor on every trial, which is globally appropriate given that participants learnt the task. However, at the beginning of blocks, participants could not know the required response yet. Furthermore, some participants failed to learn the correct response for (some of) the cues. Thus, as a more proximate measure of participants’ beliefs of what they should do, we fitted a simple Rescorla-Wagner model (Rescorla & Wagner, 1972) to the Go/ NoGo response data of each participant, simulated the action (Q) values for Go and NoGo responses on each trial, and used the difference Q_Go_ – Q_NoGo_ as a regressor to quantify the trial-by-trial relative value of making a Go relative to NoGo response. At the beginning of a block, this regressor will be zero, and it will stay (close to) zero in case participants fail to learn the correct response. We found that the more Q-values favored a Go compared to a NoGo response, the more likely were participants to first fixate the reward (Sample 1: *b* = 0.09, 95% CI [0.03 0.19], χ^2^(1) = 8.04, *p* = .005; Sample 2: *b* = 0.13, 95% CI [0.05 0.22], χ^2^(1) = 9.17, *p* = .002; Supplemental Material 4).

We furthermore performed exploratory analyses to test whether action plans affect attention beyond the first fixation, i.e., also the overall difference in dwell time to the stakes (dwell time on the reward stake minus dwell time on the punishment stake). This difference was higher when the reward stake was fixated first (Sample 1: *b* = 0.18, 95% CI [0.07 0.30], χ^2^(1) = 8.81, *p* < .001; Sample 2: *b* = 0.16, 95% CI [0.08 0.24], χ^2^(1) = 13.23, *p* < .001; not significant in either sample when only analyzing trials with both stakes fixated), showing that the first fixation anchored which stakes would receive overall more attention. Over and above this effect, action value kept shaping dwell times, such that people dwelt longer on the reward (compared to the punishment) stake for Go relative to NoGo cues (Sample 1: *b* = 0.03, 95% CI [0.01 0.05], χ^2^(1) = 4.71, *p* = .030; Sample 2: *b* = 0.03, 95% CI [0.02 0.05], χ^2^(1) = 13.79, *p* < .001; Supplemental Material 4), corroborated when approximating action plans alternatively via Q-values (Sample 1: *b* = 0.03, 95% CI [0.01 0.05], χ^2^(1) = 4.36, *p* = .037; Sample 2: *b* = 0.04, 95% CI [0.02 0.06], χ^2^(1) = 24.82, *p* < .001; Supplemental Material 4). Furthermore, dwell time was influenced by the stake magnitudes, with significantly longer dwell time on the reward stake compared to the punishment stake for more positive stakes differences (Sample 1: *b* = 0.09, 95% CI [0.05 0.13], χ^2^(1) = 16.49, *p* < .001; Sample 2: *b* = 0.12, 95% CI [0.09 0.15], χ^2^(1) = 41.59, *p* < .001; see Fig. 3B). This latter effect shows that total dwell time was not completely determined by the first fixation, which was shaped by “top down” action values, but was additionally sensitive to bottom-up saliency effects of the stake magnitudes.

In sum, we find evidence that that participants’ attention to valenced stakes information, in terms of both initial fixation and total dwell time, was synchronized to their initial action plans.

### Eye gaze predicts responses

We next assessed whether and how attention shaped the ultimate response. We used the difference in dwell times (reward minus punishment stakes) as an integral measure of total attention (Konovalov & Krajbich, 2016). We controlled for the required action to show that attention predicted the eventual response even beyond participants’ likely intentions.

The longer participants attended to rewards compared to punishments, the more likely they were to make a Go response (Sample 1: *b* = 0.13, 95% CI [0.07 0.20], χ^2^(1) = 12.20, *p* < .001; Sample 2: *b* = 0.19, 95% CI [0.13 0.26], χ^2^(1) = 28.44, *p* < .001; Fig. 3C). Furthermore, in Sample 2 (but not Sample 1), longer attention to rewards compared to punishments led to faster reaction times (Sample 1: *b* = −0.04, 95% CI [-0.09 0.02], χ^2^(1) = 1.90, *p* = .168; Sample 2: *b* = −0.03, 95% CI [-0.05 −0.01], χ^2^(1) = 4.53, *p* = .033). When considered in isolation, higher dwell time on rewards increased responding, but did not significantly affect reaction times, while higher dwell time on punishment decreased responding and slowed responses (see Supplemental Material 5). We did not observe any interaction effects between stakes and dwell time effects (see Supplemental Material 5).

As action plans both affected attention as well the ultimate response, on might wonder if the link between attention and the ultimate response was induced by action plans as a “common cause”. To exclude this possibility, all analyses using dwell times to predict responses included the required action as a regressor. Furthermore, we obtained causal evidence for an effect of attention on the ultimate response in a separate online study, in which we manipulated attention. In this study, action cues were presented simultaneously with stakes, but located in close spatial proximity to either the reward or the punishment stakes. We reasoned that the stakes closer to the action cue would receive more attention. Indeed, we observed that action cues were located closer to reward (instead of punishment) stakes resulted in more and faster Go responses. This additional dataset corroborates a causal effect of attention on the ultimate choice. For details, see the Supplemental Material 6.

In sum, we found evidence in both samples that dwell time on rewards/ punishments drove responses towards Go/ NoGo and speeded/ slowed responses, respectively, such that attention determined the eventual strength of Pavlovian biases. Tentative evidence suggested that effects of stake magnitudes and dwell times were highly similar.

### Stake magnitude and attentional effects differently relate to performance

Lastly, given that both stake magnitudes and dwell times affected responses and RTs in a highly similar way, we asked whether these effects also had similar consequences for participants’ overall performance. Crucially, stakes were controlled by the experimental protocol and were therefore unrelated to the required response on each trial. In contrast, attention was under the control of the participant. If participants fixated reward or punishment cues in line with their action goals and then let attention guide their eventual response, strong attention effects could putatively improve their performance. We performed exploratory analyses testing whether effects of stake magnitudes and dwell times on responding were related to accuracy across participants.

The effect of stake difference on responses correlated significantly negatively with accuracy, *r*(97) = −0.24, 95% CI [-0.42, −0.04], *p* = .017 (see Supplemental Material 7; after removing two outliers visible: *r*(95) = −0.26, 95% CI [-0.44, −0.06], *p* = .010; Fig. 4A), while the effect of dwell time difference correlated significantly positively with accuracy, *r*(97) = 0.45, 95% CI [0.27, 0.60], *p* < .001 (Fig. 4B). Effects were not exclusively driven by reward or punishment stakes/ dwell times, but both (in opposite directions, respectively; see Supplemental Material 4). We excluded two simpler explanations of the association between the attentional effect and task accuracy: First, this association was not driven by more accurate participants providing higher-quality eye-tracking data (see Supplemental Material 7). Furthermore, accuracy was not linked to a stronger focus on reward information (i.e., more first fixation to rewards or longer attention to rewards); if anything, more accurate participants showed a more variable gaze pattern, which support the idea that these participants could rely in their responses on their context-appropriate gaze patterns (see Supplemental Material 4).

**Figure 4.**
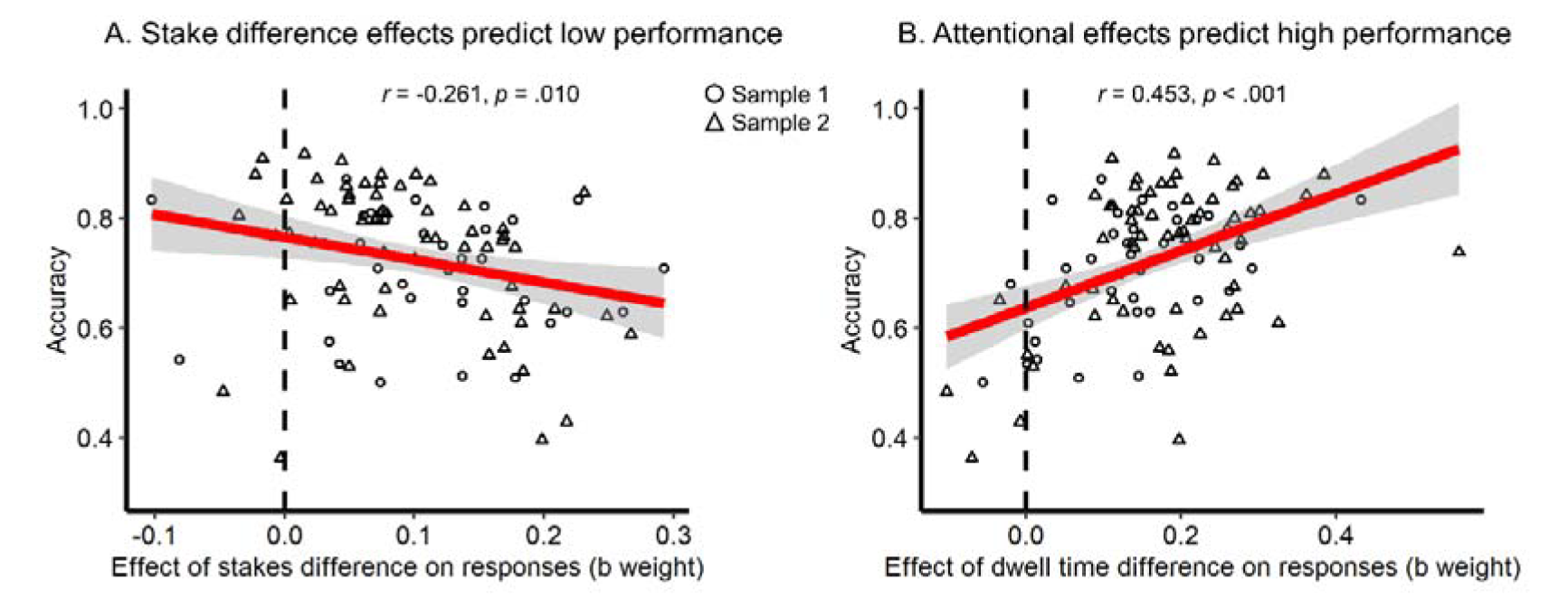
Between-subjects correlations between global Go/ NoGo task performance and stake magnitude (A) and attentional (B) effects. A. Participants with stronger effects of the stake difference on responding (i.e., steeper slopes in Fig. 2C) showed lower performance. B. Participants with stronger effects of the dwell time difference on responding (i.e., steeper slopes in Fig. 3C) showed higher performance. Individual data points reflect per-participants scores, the red line reflects the regression of accuracy on stake magnitude/ attentional effects (shades for ±1 SE). Points = individual participant effects, purple line = regression line, shading = +1 SE.

In sum, although correlational, these results suggest that strong attentional effects might facilitate performance, while strong stake magnitudes effects impair it. Based on these analyses, stake magnitude and attentional effects appear to be dissociable.

## Discussion

We report evidence from two independent samples showing that instrumental action plans steer attention towards rewards and punishments and in this way shape the input to the Pavlovian control system, triggering responses in line with those action plans. These results shed new light on the possible function of Pavlovian control. In contrast to current theories, we suggest that these biases have an important role beyond providing reasonable response defaults in novel or seemingly uncontrollable environments. Crucially, in addition, Pavlovian control can support instrumental control for efficient and robust action execution. In a novel task, participants successfully learned to perform Go and NoGo actions to various cues. Their responses and reaction times were biased by task-irrelevant information about potential reward/ punishment outcomes (stakes), similarly to previously reported Pavlovian biases. Most crucially, we found that participants aligned their attention to these stakes with their action plans: they paid more attention to reward stakes when they had to perform a Go action, and relatively more attention to punishment stakes when they had to perform a NoGo action. In turn, attention to these stakes biased ultimate responses, such that more attention to rewards increased the frequency and speed of Go responding. Exploratory between-subjects analyses showed that stronger attentional effects on choice were associated with higher performance, hinting at the adaptive nature of using attention to elicit an automatic response. In sum, these results support the notion that humans can adaptively direct attention to reward and punishment information to selectively elicit Pavlovian biases in line with their action plans.

Current theories often emphasize the “hardwired” nature of Pavlovian biases (Boureau et al., 2015; Dayan et al., 2006) that allow for fast, but inflexible responding. Under the assumption that these biases embody environmental statistics on an evolutionary time scale, they should lead to the correct response in most situations. Normative models assign a dominant role to these biases in contexts that cannot be controlled (yet) by instrumental knowledge about action-outcome relationships (Dorfman & Gershman, 2019). However, once an environment is controllable, biases should disappear. Frequent action slips reveal that Pavlovian biases continue to interfere with goal-directed behavior and require active suppression (Cavanagh et al., 2013; Swart et al., 2018). These cases of interference seem to question their putatively adaptive nature, warranting an update on previous theories.

Here, we suggest that a strong Pavlovian system can be adaptive, even in well-known environments, when it is actively brought into alignment with the goals of other (instrumental) systems. Pavlovian and instrumental control do not need to operate in a strict parallel fashion and merely interact at the output stage. Instead, we show that instrumental control can determine the input to Pavlovian control by selectively steering attention to (potentially unrelated) reward or punishment information. In this way, it sets the Pavlovian system on an “ballistic” track that will eventually lead to the intended response. Having such an auxiliary mechanism that will trigger the intended response might be particularly adaptive in real-life contexts in which the implementation of actions unfolds over time and is prone to interruption by distractors. By “aligning” Pavlovian with instrumental control, action selection becomes more robust against interference. Such an faciliatory effect of Pavlovian control is in line with our finding of better performance in participants with stronger attentional shaping of responses.

Our findings shed new light on the potential use of simple, “fast-and-frugal” systems in decision-making, motor control, and attention. These fields distinguish slow, more computationally demanding, but at the same time more flexible and “accurate” strategies against faster, less demanding, but inflexible and frequently incorrect strategies (Balleine & Dickinson, 1998; Du et al., 2022; McDougle et al., 2016; Theeuwes, 2018). The latter may yield adequate behavior only in a subset of situations, but are frequently misapplied (Beck et al., 2012; Fawcett et al., 2014; Rahnev & Denison, 2018), raising the question why they are not permanently suppressed beyond contexts of high novelty or uncertainty. In the case of Pavlovian biases, we suggest that these biases can facilitate the implementation of instrumental action plans by making them more robust against distraction. The price of infrequent motor errors caused by Pavlovian biases might be worth paying if, at the same time, the robustness of instrumental action implementation is significantly enhanced. Future research needs to address under which exact circumstances an architecture with a more flexible, sophisticated strategy and more inflexible, simple strategy warrants the infrequent errors produced by the latter.

Beyond the context of Pavlovian biases, our results extend previous literature on the upstream determinants (rather than downstream consequences) of attention allocation. Previous studies have found that, at least early in the choice process, attention appears to be randomly allocated to choice options in a way that is independent of their value (Manohar & Husain, 2013; Westbrook et al., 2020). In contrast, recent Bayesian accounts of “active sensing” have proposed that attention should be actively driven by the value and uncertainty of choice options in order to gather the maximal amount of information (Callaway et al., 2021; Jang et al., 2021; Sepulveda et al., 2020). We highlight yet another role of attention allocation: to stabilize (or even speed up) action implementation in face of delays and distraction. This role stipulates that (visual) attention is at least partly under control of ongoing motor processes—as proposed by the premotor-theory of attention (Olivers & Roelfsema, 2020; Rizzolatti et al., 1987; Sheliga et al., 1997)—as well as recent accounts highlighting that vision and visual working memory primarily serve action (Heuer et al., 2020; van Ede, 2020).

The idea of Pavlovian biases being recruited by instrumental action plans extends such accounts into the domain of value-based decision-making. It provides a potential explanation for why humans seek out a choice option right before selecting it, even when this will not reveal new information on what is the optimal choice (Hunt et al., 2016; Kaanders et al., 2021). Fixating an (appetitive) option might trigger Pavlovian biases that ensure its selection in face of distractors. Even more so, after participants have made the decision to select an option, its collection and consumption (potentially in face of competitors) might require further motor actions that can benefit from invigoration via these biases. Hence, the role of Pavlovian biases in invigorating motor programs might potentially explain phenomena of human (and animal) curiosity and information seeking (Cervera et al., 2020; Vasconcelos et al., 2015) even after the decision process is finished.

Our results also shed new light on the potential mechanisms by which attention to different choice options affects their eventual choices. Past research has not yet provided evidence on how fixating a choice option (e.g., a well-known food item like a Snickers) helps its evaluation or affords more information about it. Some accounts have proposed that value-based decisions are made by retrieving goal-relevant information or “preferences” from memory (Shadlen & Shohamy, 2016). Attention to an option could potentially facilitate the retrieval of value-related information about this option (Callaway et al., 2021). Other studies have observed effects of attention also on perceptual choices that might not require memory retrieval, suggesting that attention can also affect visual stimulus processing directly (Smith & Krajbich, 2021; Tavares et al., 2017). In contrast to all of these studies, our results suggest that attentional effects might be uncoupled from any features of the choice option and instead be “Pavlovian” in nature: attending to (any) positive information disinhibits motor cortex and facilitates selection, while attending to (any) negative information inhibits motor cortex and leads to rejection—regardless of whether this information is related to the choice option or not. Crucially, in our paradigm, positive and negative information was unrelated (and orthogonal) to the action that needed to be selected, and thus should not be incorporated into the choice process. However, even this unrelated information did bias choice. To dissociate whether attentional effects are truly driven by increased knowledge about an option’s features rather than a simple (dis-) inhibitory effect of its valence, future research should systematically manipulate the relevance of positive and negative option features to the eventual choice.

There are a few important considerations when generalizing our findings to real world situations. First, possible outcomes of a choice are often not explicitly presented to an agent. Rather, agents must make a selection among many potentially relevant pieces of information on what they deem important. Our task tried to mimic such situations by allowing agents to freely choose how much to attend to information about rewards and punishments at stake. Still, attention allocation differed from “naturalistic” free viewing settings in two important ways. Participants were not completely free to attend to the stakes, but were incentivized to do so by the secondary catch task. Furthermore, only two pieces of potential information—exemplary of positive and negative aspects of the situation—were presented, which is a drastic simplification of our information-dense environment. Future extensions of this research should provide participants with a larger set of information to select from, allowing them complete freedom to seek out any information during action preparation.

Second, in real life situations as well as in this task, people might initiate an action plan, but then change their mind. We only had access to the participants’ ultimate response, which does not allow us to disentangle situations in which they maintained a determined action plan throughout the trial from situations in which actions plans were changed based on reward/ punishment information. Neuroimaging techniques with high temporal resolution such as EEG and MEG could shed light on the dynamic interactions between motor processes and how these change as a function of attentional focus.

Third and finally, exploratory analyses suggested that participants whose ultimate response relied more strongly on attentional inputs showed higher performance. This result corroborates the postulated adaptive nature of a strong Pavlovian system that can be harnessed by instrumental systems. In contrast, the degree to which responses were shaped by the stakes magnitudes (i.e., larger magnitudes resulting in stronger Pavlovian biases) was associated with lower performance. This—at first perhaps surprising—dissociation likely arose from our task design in which stakes magnitudes were orthogonal to action requirements. When participants performed substantially above chance, stakes magnitudes had a greater potential to disturb action selection on “incongruent” trials (where The required action and the action triggered by the net stakes difference mismatched) than to facilitate it on “congruent” trials. In contrast, in many real-world contexts, it is adaptive to take into account the size of available rewards or punishments when choosing whether and how vigorously to respond.

Still, even if stakes magnitudes and attention to stakes are both meaningful contributors to choices in real-world settings, it is noteworthy that both had different consequences for performance in our task, suggestive of dissociable behavioral phenotypes. While relying on stake magnitudes might be linked to “sign-tracking” behavior previously observed in animals and humans (Flagel et al., 2009, 2010; Schad et al., 2020) and suggested to constitute a risk factor for addiction (Chen et al., 2023; Garbusow et al., 2016; Robinson & Berridge, 1993), relying on attention might be a “novel” phenotype reflecting a strategic recruitment of Pavlovian biases. To conclusively demonstrate the adaptive nature of using attention to invigorate Pavlovian biases, future studies would need to causally manipulate participants’ strategies. Such studies could for example train participants to strategically seek out reward or punishment information under a certain action plan. The ability to strategically up-or down-regulate Pavlovian biases could then be relevant for future interventions in psychopathologies characterized by aberrant biases, such as depression (Huys et al., 2016) or alcohol addiction (Chen et al., 2023; Garbusow et al., 2016; Schad et al., 2020; Sommer et al., 2017).

In sum, our results suggest a broadening of the current view of Pavlovian control: In addition to providing sensible “default” actions in novel or uncontrollable environments, a strong Pavlovian system can be adaptive even in well-known environments when its robust, almost “ballistic” nature is recruited to ensure that an action plan is implemented even in face of distraction.

## Context of this research

Much literature on Pavlovian biases has focused on situations in which these biases are maladaptive, investigating how they can be suppressed via top-down control (Cavanagh et al., 2013; Swart et al., 2018). However, stronger biases have been found predictive of better recovery from depression (Huys et al., 2016). Furthermore, initial theoretical considerations have proposed that biases could be evaded by mentally reframing a given situation (Boureau & Dayan, 2011) rather than recruiting top-down control. We pursued this line of reasoning experimentally, testing whether humans use attention to reward/ punishment cues to create a “Win”/ “Avoid” situation that helps them pursue their action goals. This perspective highlights that instrumental and Pavlovian control might more often work on concert rather than oppose each other.

## Constraints of generality

Pavlovian biases might be a universal phenomenon shared by humans and many animal species. They have been described across the animal realm, suggesting a genetic basis shared amongst humans and other animals and/or a “mandatory” acquisition very early in life in a set of diverse environments. While there are considerable individual differences in the strength of these biases (as described in this manuscript as well as previous work), the direction of their effects both within and across species is highly consistent, with reward prospect invigorating responding and threat of punishment inhibiting it. Systematically inverted biases have never been observed. In contrast, for the strategic attentional recruitment of these biases, there might be no similar “hard-wired” basis and such a strategy might be acquired by different individuals to different degrees. We speculate that, similar to the biases themselves, the direction of this strategic effect is consistent across individuals. The existence of an “inverted” strategy is highly implausible. The authors would like to highlight that the studies reported in this manuscript were conducted in English and that a significant portion of the participants were not Dutch natives (although this was not systematically assessed), suggesting that the strategic recruitment of biases occurs independently of the local culture where this research has taken place.

## Acknowledgements

**Acknowledgements:** We thank Jesse van der Spek and Helena Olraun for assistance with data collection and Peter Dayan for helpful comments and suggestions on the manuscript.

## Supplemental Material 1: Results overview full sample

Here, we report an overview over all major statistical results reported in the main text and the supplementary material. These results are based on all participants in both samples. For details on how mixed-effects regression were performed, see the Methods section of the main text.

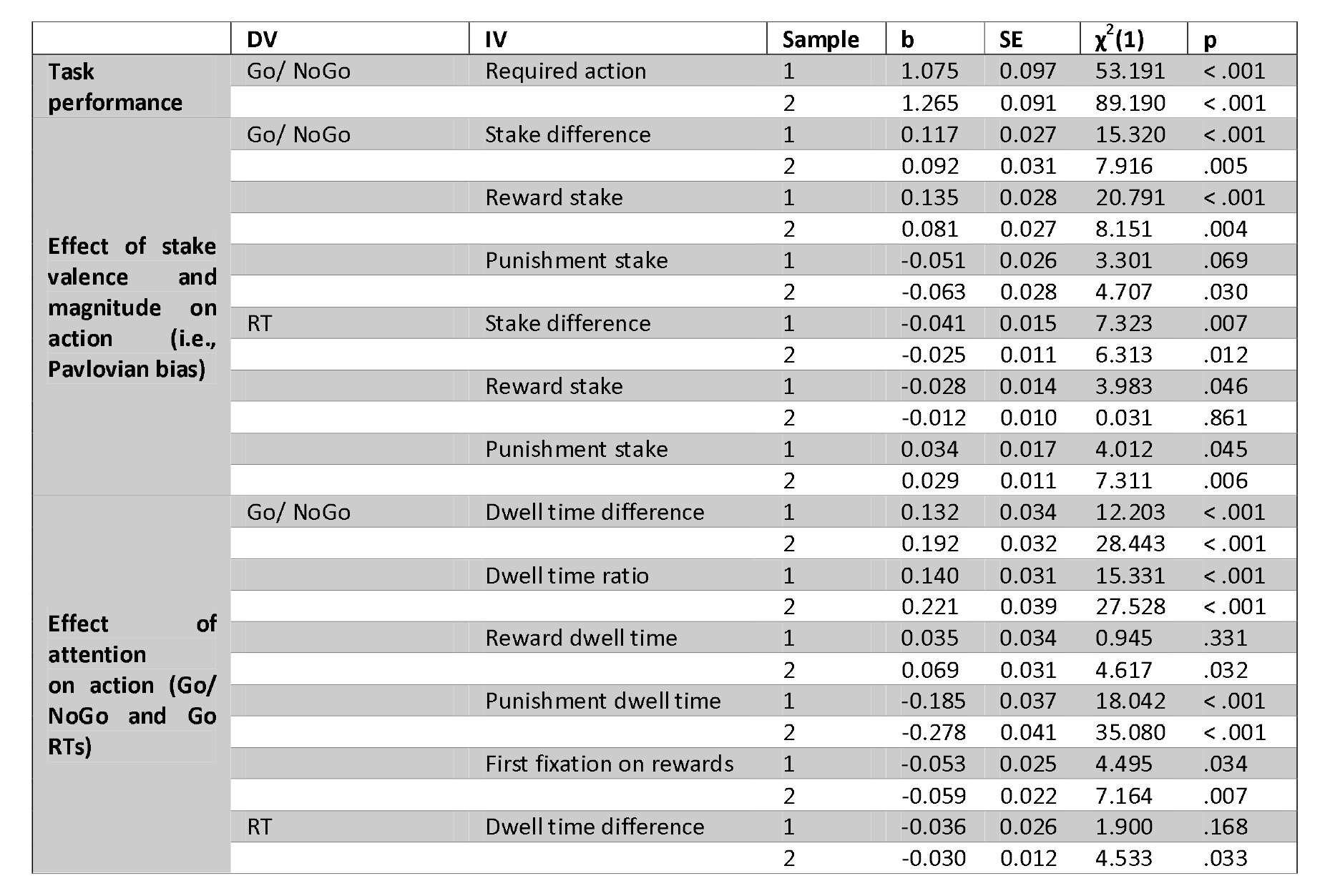

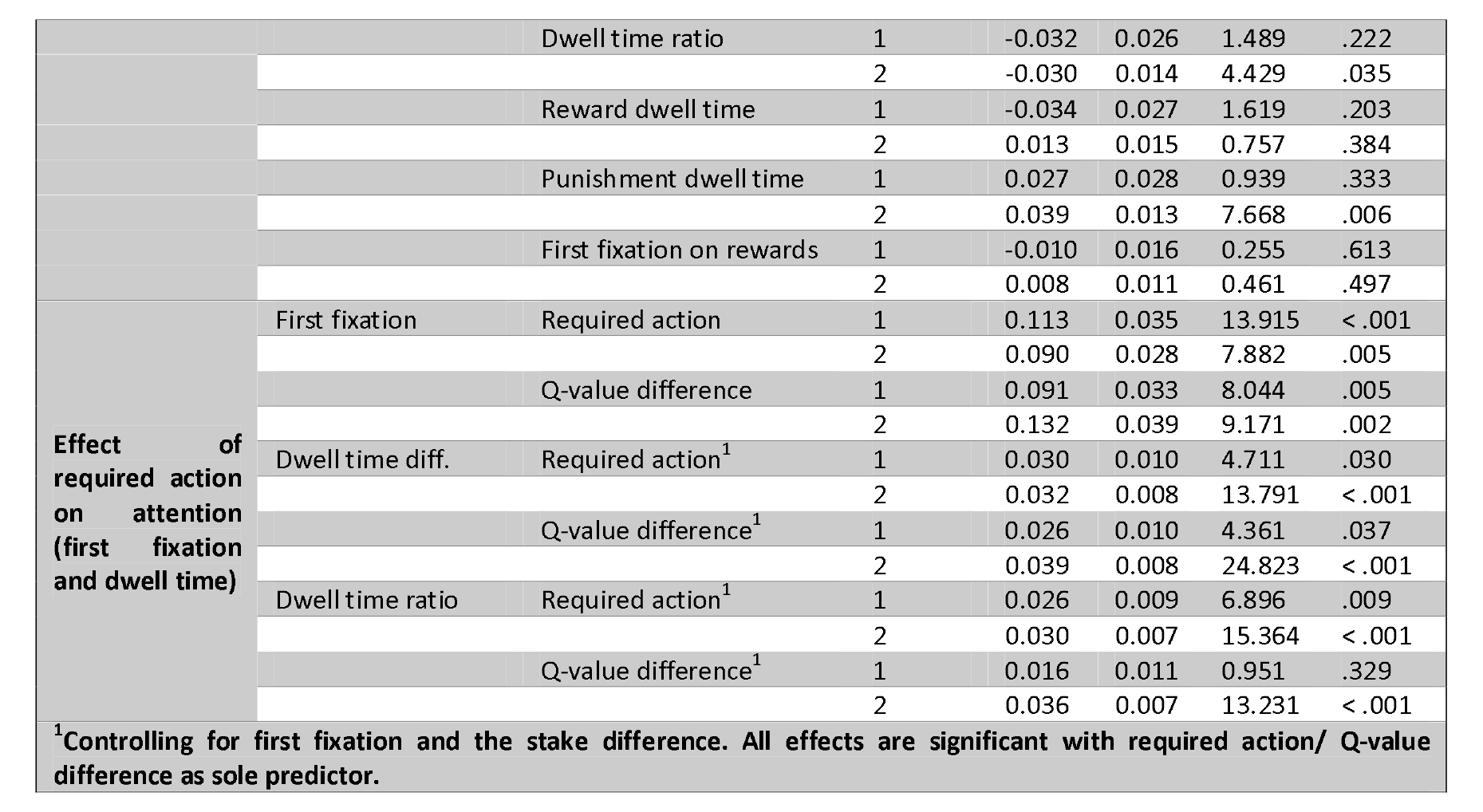

## Supplemental Material 2: Results overview: Participants not significantly above chance excluded

We report an overview over all major statistical results as reported in the main text and the supplementary material, but excluding the five (seven) participants in Sample 1 (2) that did not perform significantly above chance level, i.e., did not learn the task. For details on how mixed-effects regression were performed, see the Methods section of the main text. These analyses led to the same conclusions as the analyses based on the full samples reported in S01.

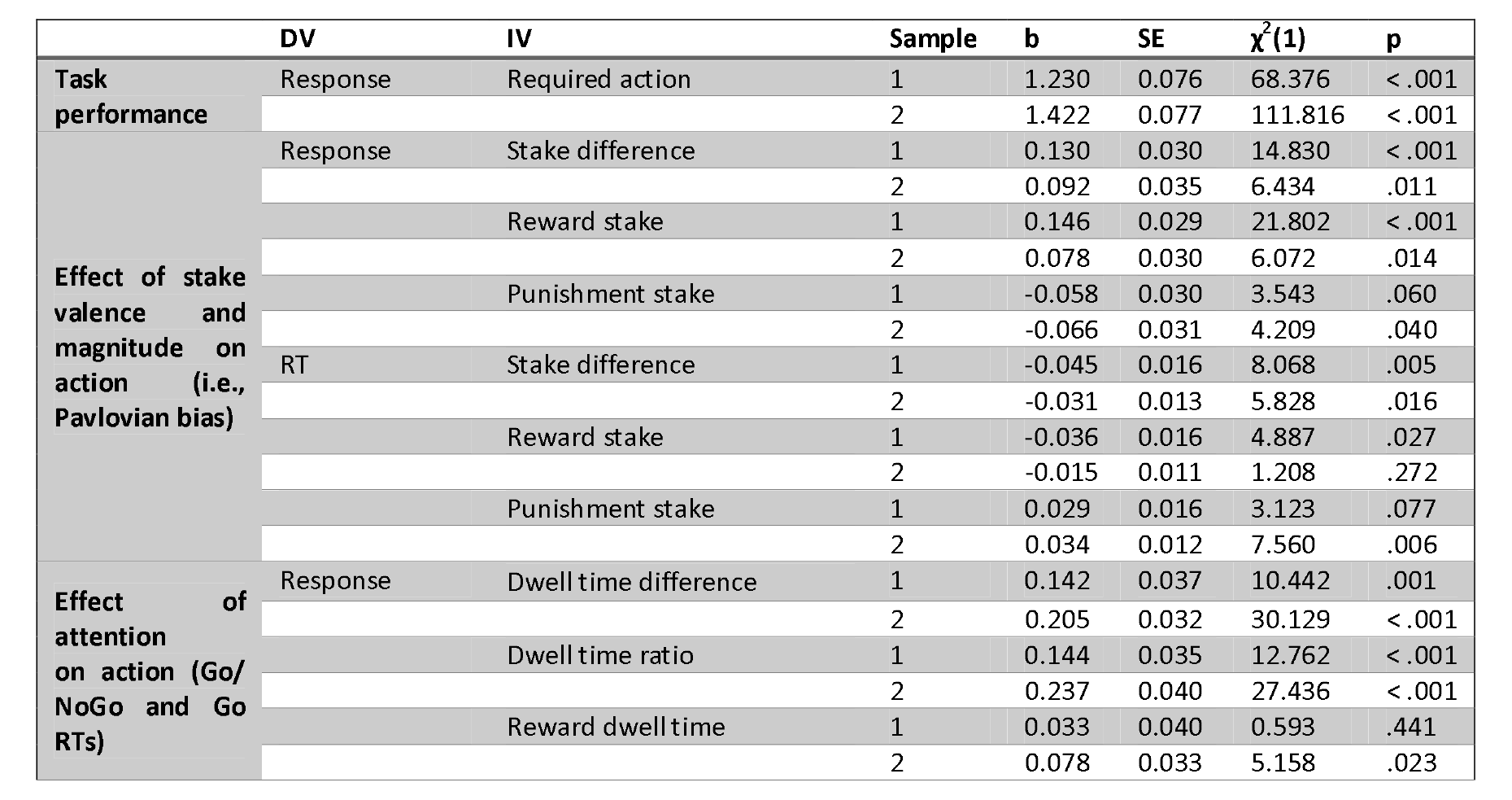

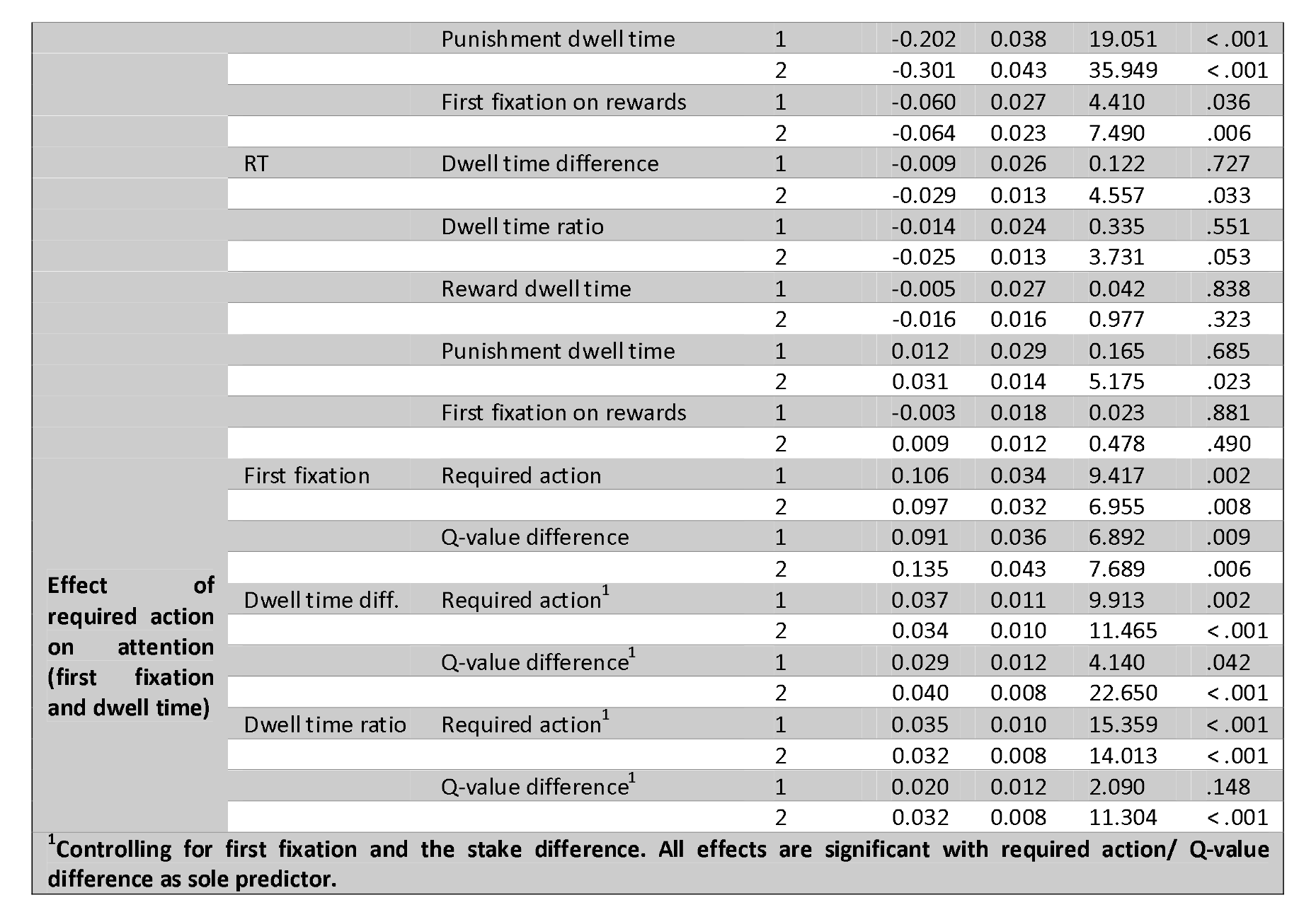

## Supplemental Material 3: Effects of stake magnitudes on responses and reaction times

Given that stake differences (reward minus punishment stake) affected both Go/ NoGo responses and reaction times, we additionally tested for separate effects of the reward and punishment stake magnitude on responses and reaction times using in mixed-effects logistic regressions (for Go/ NoGo responses) and linear regressions (for reaction times). We coded reward and punishment stake magnitudes as separate regressors (instead of as their difference).

The effect of reward stake magnitude on responses was significant in both samples (Sample 1: *b* = 0.14, 95% CI [0.08 0.19], χ^2^(1) = 20.79, *p* < .001; Sample 2: *b* = 0.08, 95% CI [0.03 0.13], χ^2^(1) = 8.15, *p* = .004; Fig. S03A), while the effect of punishment stake magnitude was only significant in Sample 2 (Sample 1: *b* = −0.05, 95% CI [-0.10 0.001], χ^2^(1) = 3.30, *p* = .069; Sample 2: *b* = −0.06, 95% CI [-0.12 −0.01], χ^2^(1) = 4.71, *p* = .030; Fig. S03B). In contrast, for RTs, higher reward stake magnitude predicted faster responses only in Sample 1 (Sample 1: *b* = −0.03, 95% CI [-0.06 −0.001], χ^2^(1) = 3.98, *p* =.046; Sample 2: *b* = −0.01, 95% CI [-0.03 0.01], χ^2^(1) = 0.03, *p* = .861; Fig. S03C), while higher punishment stake magnitude consistently predicted slower responses (Sample 1: *b* = 0.03, 95% CI [0.001 0.07], χ^2^(1) = 4.01, *p* = .045; Sample 2: *b* = 0.03, 95% CI [0.01 0.05], χ^2^(1) = 7.31, *p* = .006; Fig. S03D). Note that RTs are only available for Go responses; hence, the amount of data (and resulting statistical power) are lower compared to the Go/ NoGo response data.

In conclusion, effects of stake magnitude on driving Pavlovian biases reported in the main manuscript were driven by variations in both the reward and the punishment stake. These effects resemble effects of Pavlovian biases reported before, but in this study emerged in a graded fashion, i.e., more and faster Go responding the larger the reward stake was, and less and slower Go responding the larger the punishment stake was.

In addition, we tested whether the effect of stake difference on responses (i.e., the Pavlovian bias) became weaker over time. For this purpose, we used mixed-effects logistic regression models including stake difference, time, and their interaction. As time, we either used a) trial number across the whole task (1–264), b) trial number within each block (1–88), c) cue repetition number (1–22), or d) block number (1–3). A significant interaction would indicate that the Pavlovian bias changes over time. However, we did not find any significant interaction, neither a) for trial number across the task (Study 1: b = −0.04, 95% CI [-0.10 0.02], χ2(1) = 2.55, p = .110; Study 2: b = −0.03, 95% CI [-0.07 0.01], χ2(1) = 1.49, p = .222) nor b) for trial number within blocks (Study 1: b = −0.02, 95% CI [-0.08 0.04], χ2(1) = 0.61, p = .433; Study 2: b = −0.02, 95% CI [-0.06 0.02], χ2(1) = 1.11, p = .293), nor c) by cue repetition number (Study 1: b = −0.02, 95% CI [-0.07 0.04], χ2(1) = 0.27, p = .597; Study 2: b = −0.02, 95% CI [-0.07 0.02], χ2(1) = 0.89, p = .345), nor d) for block number (Study 1: b = −0.03, 95% CI [-0.09 0.03], χ2(1) = 1.14, p = .286; Study 2: b = −0.02, 95% CI [-0.06 0.02], χ2(1) = 0.56, p = .455). Numerically (but not significantly), the bias got weaker with time, which is to be expected given that people make less errors over time, while errors are necessary to detect the presence of a Pavlovian bias. In sum, we found no evidence that the Pavlovian bias vanishes over time.

Of note, in our pre-registration, we mentioned under “exploratory analyses” that we would fit reinforcement-learning drift diffusion models (RL-DDMs) to jointly analyze the effects of stakes/ dwell times on choices and RTs. We decided to not report the results from these models because data simulated from them was markedly different from the empirical data. We suspect that DDMs cannot capture data from this task due to i) the tight response deadline (600 ms), leading to overall fast (but regularly incorrect) responses while preventing late responses, and ii) the absence of RTs for the NoGo responses, which can be computationally dealt with, but which implies a lack of constraint on the parameters (especially the starting point bias term). Lastly, enforcing a strict response threshold is not possible in the DDM framework. Potentially, evidence accumulation frameworks in which the response thresholds decrease and eventually become zero at the response deadline might be able to accommodate such data, but likelihood functions for such models are not readily available. We encourage other researchers to reanalyze this data with more suitable modeling frameworks that might arise in the future.

**Figure SI03.**
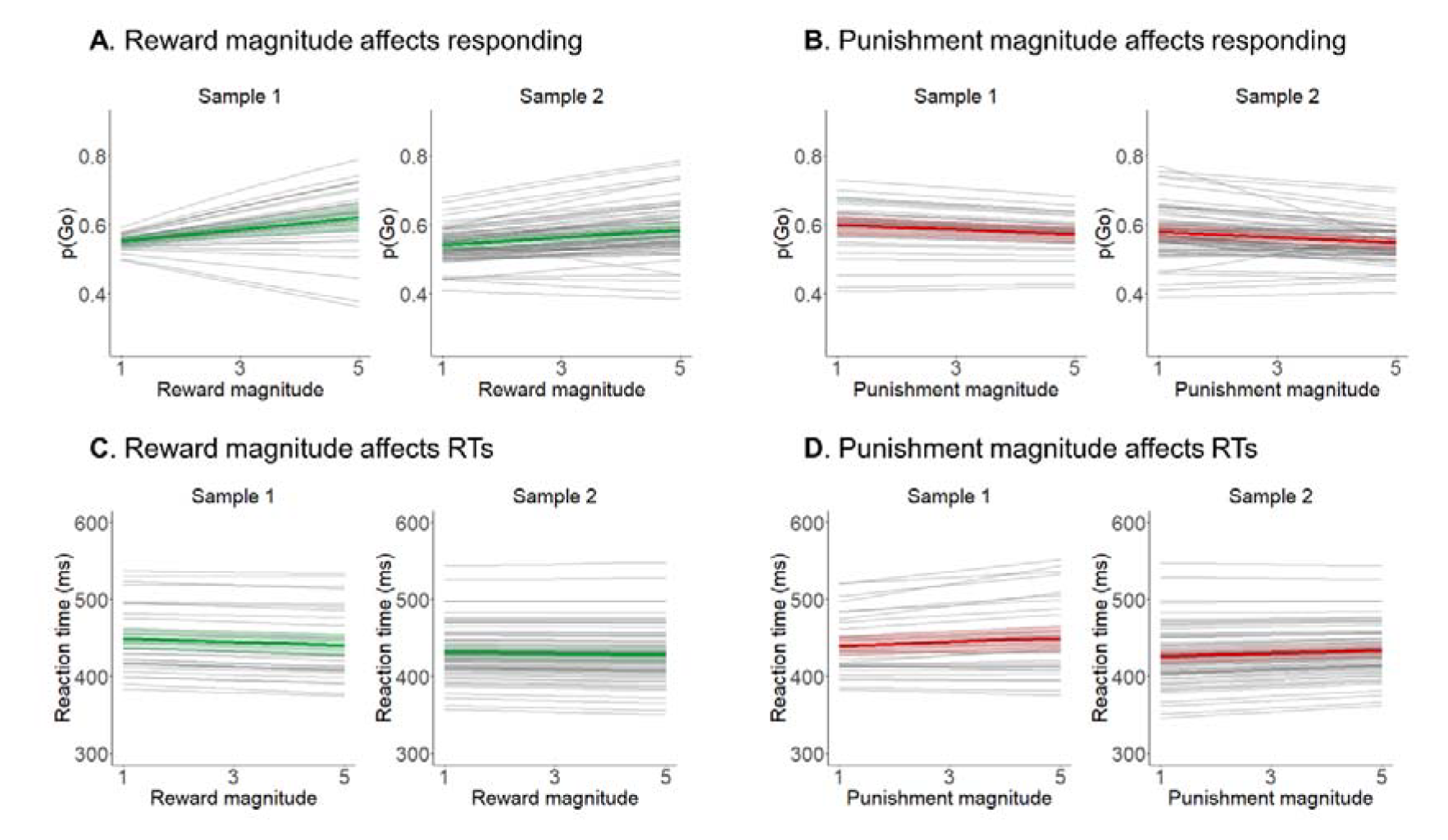
Effect of stake magnitudes on responses and reaction times. A higher reward stake magnitude led to a higher proportion of Go responses (**A**; significant in both studies), while a higher punishment stake magnitude led to a lower proportion of Go responses (**B**; only significant in Study 2). Similarly, a higher reward stake magnitude tended to speed up reaction times (**C**; significant only in Study 1), while a higher punishment stake magnitude tended to slow down reaction times (**D**; significant in both studies).

## Supplemental Material 4: Effect of action plans on attentional measures

As our first key prediction, we tested whether attention allocation to reward and punishment stake was affected by action requirements. For this purpose, we regressed attention measures (first fixation and dwell time difference) on participants’ trial-by-trial action plans (required action and Q-value differences) using mixed-effects logistic (first fixation) and linear (dwell time difference) regression. Results are reported in the main text as well as in S01. First fixations were more likely on rewards when a Go action was required/ Q-values favored Go over NoGo. Similarly, participants looked overall longer at the reward (compared to the punishment) stake when a Go action was required/ Q-values favored Go over NoGo. Taken together, all these results suggest that attention to rewards/ punishments was synchronized to participants’ action plans.

**Figure SI04.**
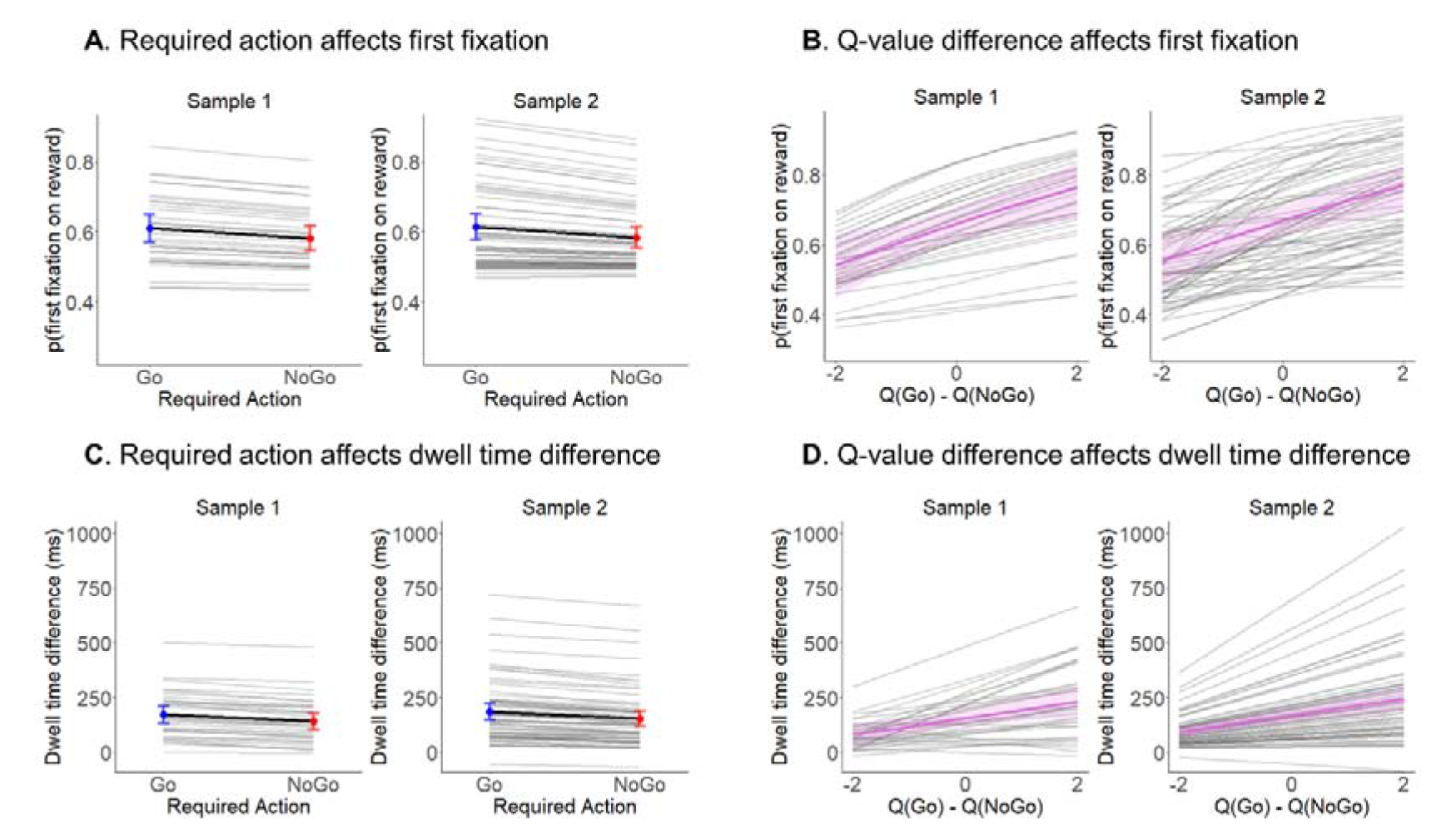
Effect of action plans on attention measures. Action requirements, i.e., whether participants should make a Go or a NoGo response based on the cue they see, biases participants’ attention during the stakes phase: A Go compared to a NoGo requirements led to a higher proportion of first fixations on the reward stake (**A**) and longer dwell time on rewards (compared to punishments) (**C**). The same finding was obtained when fitting a Rescorla-Wagner model to participants’ responses and using the Q-values based on responses from past trials to predict what participants should do on the current trial (**B** and **D**).

## Supplemental Material 5: Effect of dwell times on responses and reaction times

Given that dwell time differences (reward minus punishment dwell times) affected both Go/ NoGo responses and reaction times, we additionally tested for separate effects of reward and punishment dwell times (instead of the difference in dwell times) on responses and reaction times using in mixed-effects logistic (for Go/ NoGo responses) and linear (for reaction times) regressions. Dwell time on rewards predicted a higher proportion of Go responses significantly only in Sample 2 (Sample 1: *b* = 0.04, 95% CI [-0.03 0.10], χ^2^(1) = 0.95, *p* = .331; Sample 2: *b* = 0.07, 95% CI [0.01 0.13], χ^2^(1) = 4.62, *p* = .032; Fig. S04A). Dwell time on punishments significantly predicted a lower proportion of Go responses in both samples (Sample 1: *b* = −0.19, 95% CI [-0.26 −0.11], χ^2^(1) = 18.04, *p* < .001; Sample 2: *b* = −0.28, 95% CI [-0.36 −0.20], χ^2^(1) = 35.08, *p* < .001; Fig. S04B). Reward dwell time did not significantly predict RTs in either sample (Sample 1: *b* = −0.03, 95% CI [-0.09 0.02], χ^2^(1) = 1.62, *p* = .203; Sample 2: *b* = −0.01, 95% CI [-0.04 0.02], χ^2^(1) = 0.76, *p* = .384; Fig. S04C), but punishment dwell time predicted slower RTs in Sample 2 (Sample 1: *b* =0.03, 95% CI [-0.03 0.08], χ^2^(1) = 0.94, *p* = .333; Sample 2: *b* = 0.04, 95% CI [0.01 0.07], χ^2^(1) = 7.67, *p* = .006; Fig. S04D). Note that RTs are only available for Go responses; hence, the amount of data (and resulting statistical power) are lower compared to Go/ NoGo response data.

Interestingly, stake magnitudes and dwell times exerted highly similar effects on both responses and reaction times, with higher reward stake magnitude as well as more attention to them increased Go responding and speeded responses, while higher punishment stake magnitude as well as more attention to them decreased Go responding and slowed responses. Given that stake magnitudes and dwell times exerted such highly similar effects, one might expect them to operate through the same underlying mechanism. One consequence following from such a shared architecture is that the effects might influence each other, predicting an interaction effect. We hence performed exploratory analyses testing for such an interaction effect, reflecting whether effects of longer vs. shorter attention to the reward (punishment) stake were amplified when participants saw many vs. few potential rewards (punishments) or vice versa. The interaction between the stake difference and the dwell time difference on responses was not significant in either study (Sample 1: *b* = −0.008, 95% CI [-0.06 0.04], χ^2^(1) = 0.10, *p* = .755; Sample 2: *b* = −0.04, 95% CI [-0.08 0.002], χ^2^(1) = 3.42, *p* = .064), and neither was the case for RTs (Sample 1: *b* = 0.04, 95% CI [-0.01 0.08], χ^2^(1) = 2.25, *p* = .133; Sample 2: *b* = −0.003, 95% CI [-0.02 0.02], χ^2^(1) = 0.03, *p* = .856), providing no evidence for attention amplifying effects of stake magnitudes or vice versa.

In conclusion, longer dwell time on rewards led to more and faster responding while longer dwell time on punishments led to less and slower responding. However, effects on reaction times were only significant in the punishment domain. We did not find evidence for an interaction between stake magnitudes and dwell times, yielding no conclusive evidence whether both effects rely on the same underlying mechanism or not.

**Figure SI05.**
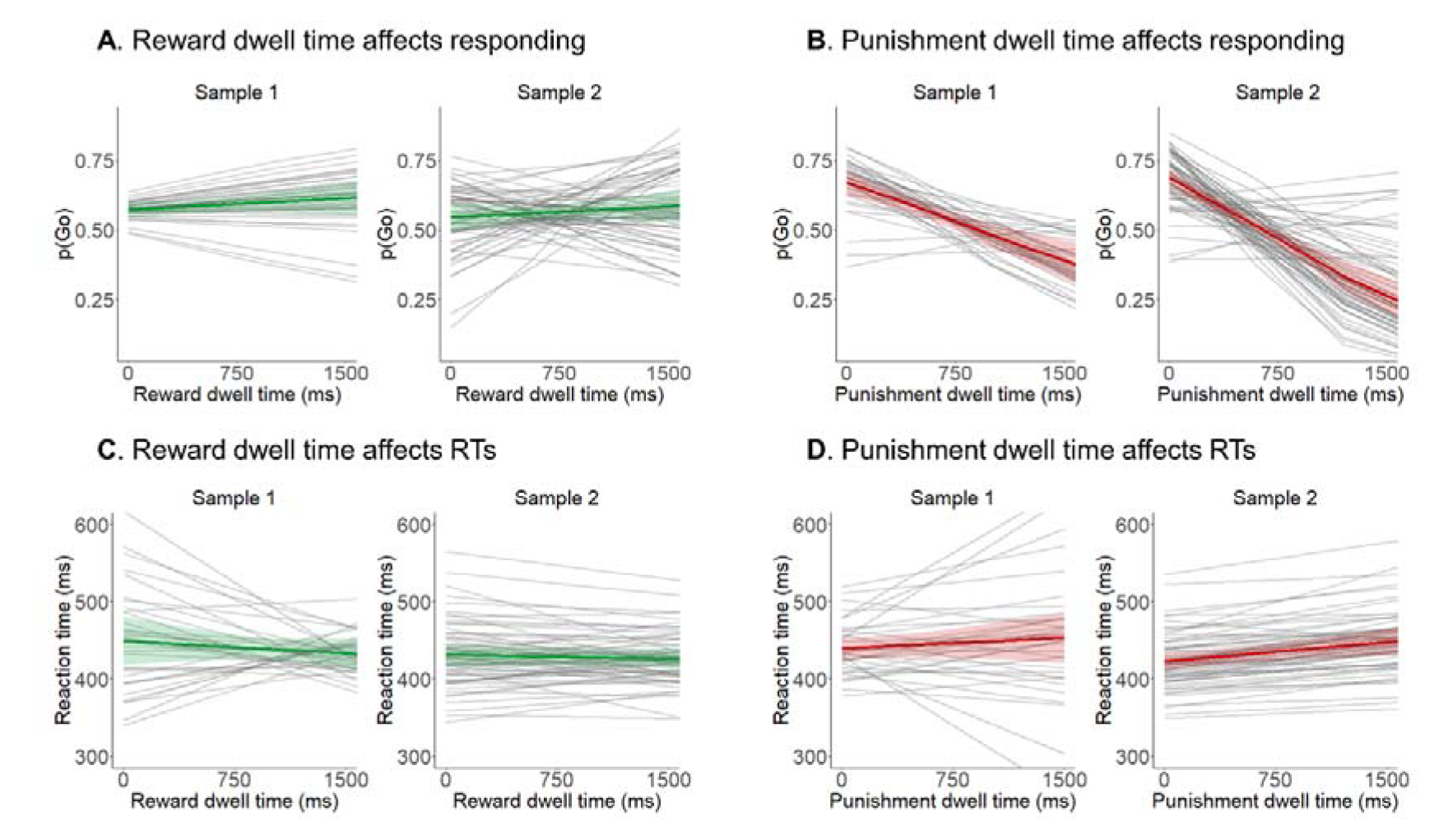
Effect of dwell times on responses and reaction times. Higher absolute dwell time on rewards led to a higher proportion of Go responses (**A**; only significant in Study 2), while higher absolute dwell time on punishments led to a lower proportion of Go responses (**B**; significant in both studies). Similarly, higher dwell time on rewards tended to speed up reaction times (**C**; though not significant in either study), and higher dwell time on punishment tended to slow down reaction times (**D**; only significant in Study 2).

## Supplemental Material 6: Supplementary online study manipulating attention to reward and punishment stakes

In the results from our eye-tracking studies reported in the main text, we observed an effect of (manipulated) action requirements on eye-gaze (first fixation and dwell time) and an effect of (measured) eye-gaze on the ultimate response. Given that both action requirements and eye-gaze predicted the ultimate response, on might wonder whether the link between eye-gaze and the ultimate response was spurious, induced by action plans as a “common cause” (an instance of the “third variable problem”). Note that all analyses regressing responses onto dwell time reported in the main text controlled for the action plans. In addition, we tested for a causal effect of attention to reward/ punishment information on responses in a separate online study in which we manipulated attention. This study was performed as a thesis project for Bachelor students at the beginning of the COVID-19 pandemic.

### Participants and Exclusion Criteria

We collected data from 34 participants (M_age_ = 22.4, SD_age_ = 2.1, range 19–27; 18 female, 31 right-handed). Data collection and analyses were pre-registered (https://osf.io/kzdhm). Data was collected under a stopping rule of N = 55 as maximal sample size or May 10, 2020 as final data collection date (set by financial/ time constraints). As pre-registered, we conducted all analyses in two ways, once including all participants and once excluding participants who a) guessed the research hypotheses (zero participants) or b) did not significantly perform above chance (based on a per-participant logistic regression with response as dependent and required action as independent variable, with p < .05 as cut-off; three participants). Both ways led to identical conclusions.

We recruited participants via the SONA Radboud Research Participation System of Radboud University. Participants were required to be at least 18 years old, understand English at a sufficient level (self-reported), not be color-blind, perform the experiment on a PC with a keyboard (no phones or tablets) and complete the study within a maximum of 90 minutes (i.e., 1.5 times the expected completion time). The experiment was administered via the Gorilla platform (Anwyl-Irvine, Dalmaijer, Hodges, & Evershed, 2020). After providing informed consent and demographic information on age, gender, and handedness, participants completed the “reversed-dot-probe” version of the Motivational Go/ NoGo Task for 30-40 minutes (see below). Afterwards, they filled out the brief (13-item) version of the Self-Control Scale (SCS) (Tangney, Baumeister, & Boone, 2004) and the Behavioral Activation/ Behavioral Inhibition System Scales (BIS/BAS) (Carver & White, 1994). Additionally, participants completed two vignettes (measuring omission bias) in which they rated the experienced regret and responsibility of two football coaches who won/ lost a match, afterwards changed/ kept their match plan, and then lost the next game (adapted from (Zeelenberg, van den Bos, van Dijk, & Pieters, 2002). Finally, participants performed a debriefing questionnaire asking them to a) guess the hypotheses of the experiment, b) report any (non-instructed) strategies they used, and c) guess whether the additional instructions helped them perform the task better. Participants were then debriefed about the purposes of the study. In compensation for participation, participants received 1 hour of course credit. Furthermore, participants with at least 60% accuracy in the Go/ NoGo task received tickets (proportional to performance) for a lottery featuring two 20€ gift card vouchers. Research was approved by the local ethics committee of the Faculty of Social Sciences at Radboud University (proposal no. ECSW-2018-171).

### Task

Participants performed an adapted version of the Motivational Go/ NoGo learning task termed “reverse-dot-probe version” (Fig. S06A). On each trial, they first saw how many points they could win for a correct response (printed in green font with a “+”) or lose for an incorrect response (printed in red font with a “-”, termed “stakes”). Stakes varied between 10 and 90 points drawn from a uniform distribution. Reward and punishment stake were presented on the left/ right side of the screen, with positions counterbalanced across blocks. Participants were instructed to attend to the stakes because these were relevant for a catch task implemented on some of the trials (see below). After 500 ms, in addition to the stakes, one out of four action cues (letter from the Agathodaimon alphabet) appeared on the screen, which required either a Go response (space bar press) or a NoGo response (no button press). Participants had to learn the correct response from trial-and-error and respond within 1,500 ms. The action cue was presented in close proximity to either the reward stake or the punishment stake, nudging participants to direct more attention to one of the two stakes. Cue position was counterbalanced across trials and orthogonal to action requirements. After a brief fixation cross screen (700 ms), participants received the outcome (either the reward or the punishment stake previously shown) displayed for 1,500 ms. Feedback was probabilistic in that 86% (12 out of 14) trials were “valid” with a correct response winning points and an incorrect response losing points, while the remaining 14% of trials were “invalid” with a correct response losing points and an incorrect response winning points. Trials ended with a variable inter-trial interval (uniform distribution from 1,100 ms till 1,900 ms in steps of 100 ms).

On 12 trials within the first two blocks, after the outcome phase, a catch task occurred. Reward and punishment stake magnitudes were presented together with a “decoy” number (all numbers printed in white font on black boxes without +/- signs, random assignment of numbers to positions). Participants had to indicate the “other” outcome they could have received (i.e., points-to-be-won in case they lost points, points-to-be-lost in case they won points) by clicking on it with the mouse within 20 seconds. The catch task required participants to memorize the exact stake magnitudes seen earlier in the trial, incentivizing attention to them. For the latter two blocks, we did not include any catch trials to not interfere with participants applying the additional instructions (see below).

After the second block, participants received additional instructions that explicitly encouraged them to look at the reward stake in case they planned to perform a Go response, and look at the punishment stake in case they planned to perform a NoGo response. In this way, we aimed to test whether participants could voluntarily align their attention with their action plans and in this way reduce the effect of the action cue’s position on responses.

Participants completed 224 trials split into four blocks à 56 trials, each blocks featuring four novel cues with 14 repetitions. Trial features (action cue identity, action requirement, stake magnitudes and positions, ITI) were controlled by one of ten pseudo-randomly drawn “spreadsheets” (preventing cue to repeated on more than two consecutive trials) randomly allocated to participants.

### Data Preprocessing

In line with the pre-registration, we excluded reaction times shorter than 300 ms from all analyses (as those are too fast to be induced by the presented cue). Using 200 ms as alternative cut-off (as used in our eye-tracking samples) did not change the conclusions.

### Analyses

We analyzed participants’ responses (Go/ NoGo) using mixed-effects logistic regression models and their reaction times using mixed-effects linear regression as implemented in the lme4 package in R (Bates, Mächler, Bolker, & Walker, 2015). For all categorical independent variables, sum-to-zero coding was used. Continuous dependent and independent variables were standardized such that regression weights can be interpreted as standardized regression coefficients. We included all possible random intercepts, slopes, and correlations to achieve a maximal random effects structure (Barr, Levy, Scheepers, & Tily, 2013). *P*-values were computed using likelihood ratio tests with the package afex (Singmann, Bolker, Westfall, & Aust, 2018). We considered *p*-values smaller than α = 0.05 as statistically significant.

As pre-registered (https://osf.io/kzdhm), firstly, we tested whether the action cue position (i.e., the cue being closer to the reward stake or to the punishment stake) as a proxy for participants’ induced attention affect their Go/ NoGo responses and reaction times, expecting a main effect of cue position. Secondly, we tested whether instructing people to attend to stake that matched their action plan reduced the effect of cue position, expecting an interaction between cue position and instructions. We tested both hypotheses in a single model (a logistic regression model for responses, a linear regression model for reaction times) featuring the regressors required response (Go/ NoGo), cue position (on the reward/ punishment side), and instructions (before /after) as well as all possible interactions. As mentioned in the pre-registration, we also report the interaction between required action and instructions as well as the three-way interaction between required action, cue position, and instructions.

Furthermore, we specified two exploratory analyses in our pre-registration. Firstly, we tested whether the difference in stakes (reward minus punishment stake) affected participants’ responses and reaction times, expecting more positive differences to lead to more and faster Go responses. For this purpose, we fitted a model with stake difference as sole regressor. Secondly, we calculated participants’ mean score on the self-control scale (SCS), BIS and BAS scales and regret judgements and tested whether these scores modulated participants’ cue position effect. For this purpose, we fitted a new model for each score featuring cue position, the respective score, and their interaction.

**Figure SI06.**
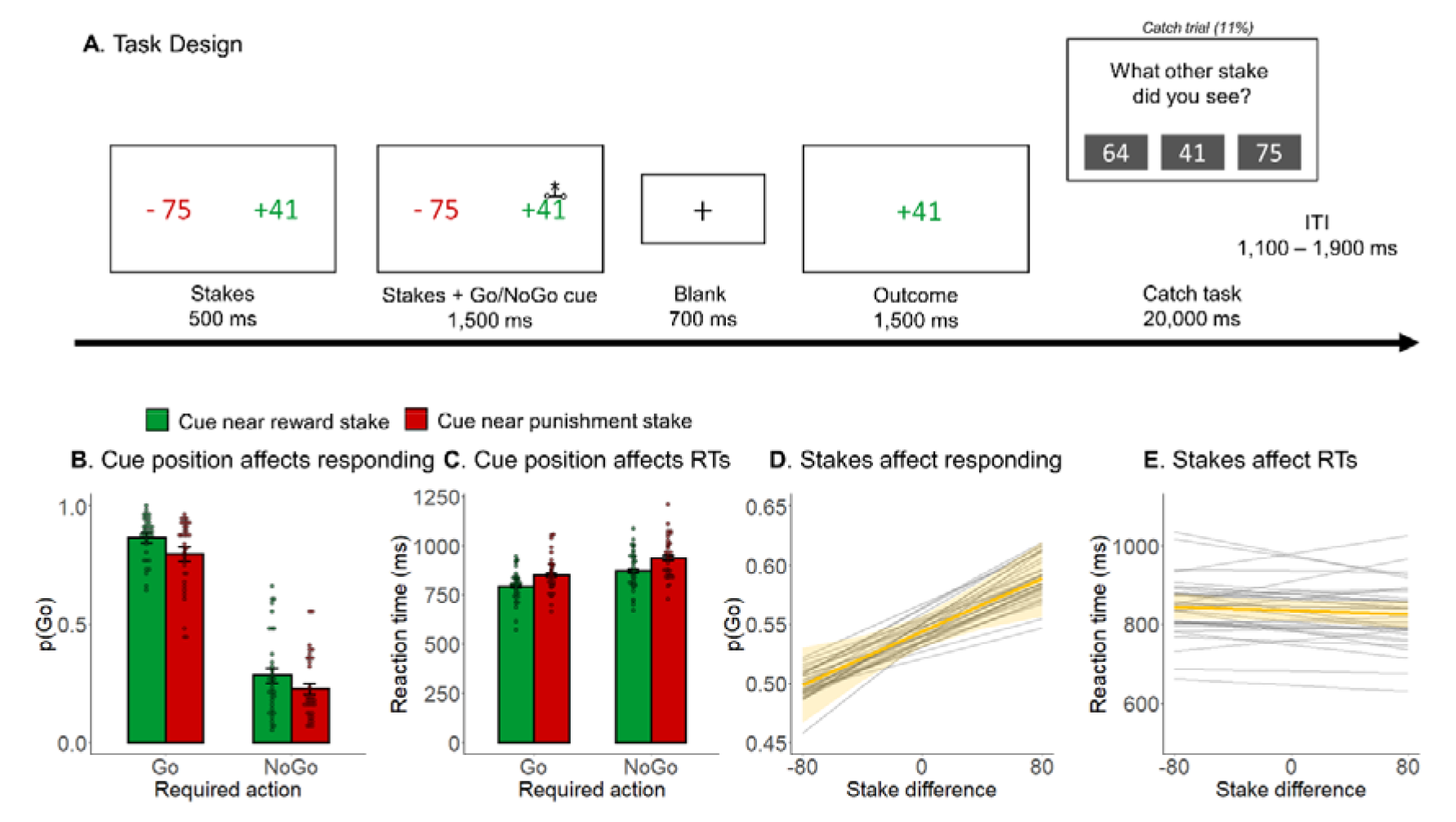
Task design and results from the online study manipulation attention to reward and punishment information **A.** Task design. On each trial, participants saw many points they could win for correct responses or lose for incorrect responses (“stakes”). After 500 ms, a Go/ NoGo action cue was displayed either next to the reward or the punishment stake, nudging participants to direct more attention to the respective stake. Participants learned whether a cue required a Go or NoGo response from trial-and-error. Outcomes are delivered in a probabilistic manner (86% feedback validity). On catch trials, participants indicated which other stake (i.e., the one they did not receive as an outcome) they had seen before. **B.** Proportion of Go responses as a function of action requirement and cue position. Participants performed significantly more Go responses to Go cues than NoGo cues and when cues were presented next to the reward stake compared to the punishment stake. **C.** Reaction times as a function of action requirement and cue position. Participants showed significantly faster responses to Go cues than NoGo cues and when cues were presented next to the reward stake compared to the punishment stake. **D.** Proportion of Go responses as a function of stake difference (reward minus punishment stake). As net stakes became more positive, participants performed significantly more Go responses. **E.** Reaction times as a function of stake difference (reward minus punishment stake). As net stakes became more positive, participants became faster, but this effect was not significant.

### Results

Overall, participants learned the Go/ NoGo task (% correct: *M* = 79.0, *SD* = 12.0, range 52.7– 94.2), performing significantly more Go responses to Go cues than NoGo cues (main effect of required action: *b* = 1.60, 95% CI [1.33 1.88], χ^2^(1) = 54.53, *p* < .001). Three participants did not perform significantly above chance (per-participant logistic regression with response as dependent and required response as independent variable, which is significant for accuracy levels of at least 56%). In line with our pre-registration, we report results with and without these participants. Performance in the catch task was above chance (3 response options imply a chance level of 33.3%; a one-sided binomial test based on 12 trials is significant for 63% accuracy and higher) in only 25 out of 34 participants. Also, the group-level performance was hardly above chance (% correct: *M* = 66.4, *SD* = 18.6, range 25.0–81.7), likely reflecting that this task was very demanding.

Firstly, in line with our pre-registration, we tested whether the cue position (action cue on the reward/ punishment side) affected participants’ Go/ NoGo responses. Participants performed more Go responses when the action cue was on the side of the reward stake compared to the side of the punishment stake (main effect of cue position: *b* = 0.19, 95% CI [0.09 0.29], χ^2^(1) = 10.90, *p* < .001; Fig. S06B), suggesting that increased attention to rewards (compared to punishments) induced more Go responses. Similarly, participants performed faster Go responses when the action cue was on the side of the reward stake compared to the side of the punishment stake (main effect of cue position: *b* = −0.03, 95% CI [-0.04 −0.02], χ^2^(1) = 25.70, *p* < .001; Fig. S06C). These results suggested that more attention directed to reward/ punishment stake causally affects participants’ responses and reaction times in the fashion of Pavlovian biases.

Secondly, in line with our pre-registration, we tested whether the effect of cue position became smaller after participants were instructed to attend to the stake that matched their action plan. The interaction effect between cue position and instructions was not significant (*b* = −0.03, 95% CI [-0.10 0.04], χ^2^(1) = 0.55, *p* = .458), providing no evidence for responses becoming less affected by the cue position once participants tried to voluntarily deploy their attention. In fact, the sign of the effect suggested the effect of cue position to become stronger (instead of weaker) after additional instructions were administered. However, there was a significant interaction between required action and instructions (*b* = −0.38, 95% CI [-0.50 −0.25], χ^2^(1) = 29.28, *p* < .001), suggesting that participant overall performed better after receiving instructions. In absence of a control group, this effect cannot be disentangled from an increase in performance over time, providing inconclusive evidence for whether instructions affected participants’ responses or not. The three-way interaction effect between required action, cue position, and instruction was not significant (*b* = 0.01, 95% CI [-0.06 0.07], χ^2^(1) = 1.78, *p* = .182). Apart from responses, also the effect of cue position on reaction times was not significantly changed by instructions (*b* = 0.01, 95% CI [-0.003 0.02], χ^2^(1) = 1.65, *p* = .199), and neither was the interaction between required action and instructions (*b* = 0.001, 95% CI [-0.01 0.01], χ^2^(1) = 0.04, *p* = .840) nor the three-way interaction effect between required action, cue position, and instruction (*b* = −0.0003, 95% CI [-0.01 0.01], χ^2^(1) = 0.004, *p* = .948) significant.

Thirdly, as part of the exploratory analyses mentioned in the pre-registration, we tested whether the difference in stakes (reward minus punishment stake) affected participants’ responses or reaction times. As expected, as the difference in stakes increased (relatively more points to win than to lose), participants performed significantly more Go responses (*b* = 0.08, 95% CI [0.03 0.12], χ^2^(1) = 8.15, *p* = .004; Fig. S06D), suggesting that the difference in available rewards/ punishments biased their responses in the fashion of Pavlovian biases. Reaction times were not significantly affected by the stake difference (*b* = −0.004, 95% CI [-0.01 0.004], χ^2^(1) = 1.01, *p* = .316; Fig. S06E).

Fourthly, as part of the exploratory analyses mentioned in the pre-registration, we tested whether the effect of cue position on responses was predicted by participants’ score on the self-control scale (SCS), the BIS and BAS scales, or the regret and responsibility ratings in the omission bias vignettes. We did not find any significant modulation of the cue position effect by SCS scores (*b* = - 0.03, 95% CI [-0.09 0.06], χ^2^(1) = 0.70, *p* = .403), BAS Drive scores (*b* = −0.04, 95% CI [-0.11 0.03], χ^2^(1) = 1.03, *p* = .310), BAS Reward Responsiveness scores (*b* = −0.01, 95% CI [-0.08 0.05], χ^2^(1) = 0.10, *p* = .756), rated regret for changing the match plan after a previous football win (*b* = −0.02, 95% CI [-0.10 0.07], χ^2^(1) = 0.14, *p* = .710), rated responsibility asymmetry when changing/ keeping the match plan after a previous football win (*b* = 0.02, 95% CI [-0.04 0.08], χ^2^(1) = 0.39, *p* = .532), rated regret for changing the match plan after a previous football defeat (*b* = −0.01, 95% CI [-0.07 0.05], χ^2^(1) = 0.10, *p* = .750), or rated responsibility asymmetry when changing/ keeping the match plan after a previous football defeat (*b* = −0.004, 95% CI [-0.09 0.08], χ^2^(1) = 0.01, *p* = .933). However, the cue position effect was significantly modulated by BIS scores (*b* = −0.07, 95% CI [-0.13 −0.01], χ^2^(1) = 4.32, *p* = .038) with participants with higher BIS scores showing weaker cue position effects, and by BAS Fun Seeking scores (*b* = −0.07, 95% CI [-0.14 −0.01], χ^2^(1) = 4.64, *p* = .031) with participants with higher BAS scores showing again weaker cue position effects. Given the sample only comprised 34 participants and several between-participants analyses were run, these results should be interpreted with caution.

We repeated all analyses while excluding three participants who did not perform significantly above chance in the Go/ NoGo task. Firstly, still, participants performed more (*b* = 0.18, 95% CI [0.08 0.29], χ^2^(1) = 10.13, *p* = .001) and faster (*b* = −0.03, 95% CI [-0.05 −0.02], χ^2^(1) = 26.84, *p* < .001) Go responses when the action cue was on the side of the reward stake compared to side of the punishment stake. Secondly, the effect of cue position on responses was again not significantly different after compared to before additional instructions were administered (*b* = −0.02, 95% CI [-0.10 0.06], χ^2^(1) = 0.24, *p* = .623), but the effect of required action was again stronger after compared to before responses (*b* = −0.41, 95% CI [-0.55 −0.27], χ^2^(1) = 23.39, *p* < .001), with again no significant three-way interaction (*b* = 0.01, 95% CI [-0.07 0.09], χ^2^(1) = 0.06, *p* = .800). Regarding reaction times, again, neither the effect of cue position (*b* = 0.01, 95% CI [-0.003 0.02], χ^2^(1) = 1.98, *p* = .159) nor the effect of required action (*b* = 0.003, 95% CI [-0.01 0.02], χ^2^(1) = 0.28, *p* = .597) was significantly modulated by instructions, and neither was the three-way interaction significant (*b* = - 0.001, 95% CI [-0.01 0.01], χ^2^(1) = 0.08, *p* = .779). Thirdly, as the stake difference increased, participants again performed significantly more Go responses (*b* = 0.06, 95% CI [0.01 0.11], χ^2^(1) = 5.72, *p* = .017), but not significantly faster responses (*b* = −0.006, 95% CI [-0.01 0.002], χ^2^(1) = 2.33, *p* = .127). Fourthly, we again did not find any significant modulation of the cue position effect by SCS scores (*b* = −0.04, 95% CI [-0.11 0.03], χ^2^(1) = 1.25, *p* = .264), BAS Drive scores (*b* = −0.02, 95% CI [-0.09 0.05], χ^2^(1) = 0.30, *p* = .582), BAS Reward Responsiveness scores (*b* = −0.01, 95% CI [-0.08 0.05], χ^2^(1) = 0.10, *p* = .751), rated regret for changing the match plan after a previous football win (*b* = 0.02, 95% CI [-0.07 0.09], χ^2^(1) = 0.27, *p* = .603), rated responsibility asymmetry when changing/ keeping the match plan after a previous football win (*b* = 0.02, 95% CI [-0.04 0.09], χ^2^(1) = 0.23, *p* = .632), rated regret for changing the match plan after a previous football defeat (*b* = −0.004, 95% CI [-0.07 0.06], χ^2^(1) = 0.01, *p* = .909), or rated responsibility asymmetry when changing/ keeping the match plan after a previous football defeat (*b* = 0.007, 95% CI [-0.08 0.10], χ^2^(1) = 0.02, *p* = .877). The modulation by BIS scores was not significant any more (*b* = −0.06, 95% CI [-0.13 0.004], χ^2^(1) = 2.33, *p* = .127), while the modulation by BAS Fun Seeking scores was still significant (*b* = −0.06, 95% CI [-0.13 −0.003], χ^2^(1) = 4.19, *p* = .041). Overall, analyses excluding the three participants who did not perform the Go/ NoGo task significantly above chance led to identical conclusions as analyses including all participants.

### Discussion

In this study, we manipulated attention by displaying Go/ NoGo action cues next to either the reward or punishment stake, nudging participants to pay relatively more attention to the stake that we next to the action cue. We obtained causal evidence that attention to reward information (compared to punishment information) leads to more Go (compared to NoGo) responses as well as to faster responses. We did not find evidence for instructions to voluntarily deploy attention in line action plans reducing the attentional effect. Potentially, the task was too demanding and the trial time course too fast for participants to voluntarily steer attention in a way that supported their action plans. Future studies might use different instructions or an altered task design that gives participants more time to deploy attention before they perform an action.

Furthermore, we found evidence for overall stake differences (reward minus punishment stake) biasing responses (but not reaction times) in the fashion of Pavlovian biases. These results support the effect of stake differences on responses reported in the main text. Finally, we did not find any strong modulation of the attentional effect by self-reported measures such as the Self-Control Scale, the BIS/ BAS scales, or regret and responsibility ratings in two vignettes measuring omission biases. Although there was some evidence for stronger BIS and BAS Fun Seeking scores predicting weaker attention effects, these results should be treated with caution given the limited sample size and the higher number of tests. Future studies should test for such links in larger samples. In sum, the core conclusion is that the results of this study support a causal effect of attention on Go/ NoGo responses.

## Supplementary Material 7: Effects of stake magnitudes and dwell times on responses predict interindividual differences in task performance

Both stakes and dwell times affected Go/ NoGo responses (and reaction times) in a similar way, i.e., a higher reward stake as well as more attention to it increased Go responding and speeded responses, while a higher punishment stake as well as more attention to it decreased Go responding and slowed responses. Given such highly similar effects, one might expect them to operate through the same underlying mechanism. First, one consequence following from such a shared architecture is that effects should influence each other, i.e., the presence of a higher stake could alter the impact of dwell times on responses, or vice versa, which predicts an interaction effect. However, we observed no evidence for such an interaction effect (see S05), tentatively suggesting that effects operate independently of each other (though curiously with highly similar consequences).

An alternative way of assessing how comparable these effects are is to probe their consequences for task performance across participants: Does letting responses be strongly guided by stake differences (reward minus punishment stake magnitudes) vs. strongly guided by dwell time differences (reward minus punishment dwell times) have similar or different consequences for overall performance in the Go/ NoGo task? For this purpose, we re-fitted regression models across both samples, extracted per-participant regression coefficients (fixed-effect plus participant-specific random effect), and correlated these coefficients with participant overall performance (% correct responses).

Performance was significantly lower in those participants in which stake difference more strongly shaped their responses (Figure S08A, B). This finding was in stark contrast to significantly higher performance in those participants in which dwell time differences (reward minus punishment dwell time) more strongly affected response. It is noteworthy that the stake differences are experimentally controlled, and thus purely “bottom-up”, while in contrast, dwell time differences were under participants’ control and synchronized to action plans, both directly (effect on dwell time difference) and indirectly (effect on first fixations).

We performed control analyses to exclude the possibility that the association between attentional effects on responses and task performance was driven by better performing participants showing higher eye-tracking data quality. First, we computed the number of trials with any (opposed to no) fixation on any of the two stakes. This number was significantly positively correlated with performance, *r*(97) = 0.23, 95% CI [0.03, 0.41], *p* = .025, but not with the attentional effect on responses, *r*(97) = 0.13, 95% CI [-0.07, 0.32], *p* = .208. When using both task performance and number of trials with any fixation to predict attention effects in a multiple linear regression, the effect of task performance was still strongly significant, *t*(96) = 4.79, *p* < .001. Second, we calculated the total time (in ms) that people attended to any of the two stakes objects. This number was neither significantly correlated with performance, *r*(97) = 0.09, 95% CI [-0.11, 0.28], *p* = .389, nor with the attentional effect on responses, *r*(97) = 0.13, 95% CI [-0.07, 0.32], *p* = .183, and when using both task performance and total fixation time to predict attention effects in a multiple linear regression, the effect of task performance was still strongly significant, *t*(96) = 4.90, *p* < .001. In sum, it is unlikely that the correlation between performance and attentional effects on responses is driven by more accurate participants providing higher-quality eye-tracking data.

Furthermore, we performed control analyses checking whether performance, being associated with how many rewards (rather than punishments) participants received, was associated with differential fixation patterns (more first fixations or longer fixations) to reward vs. punishment stakes. It is possible that performance affects information search: high performing participants can reasonably expect to receive rewards most of the time, so they might be more interested in and attend more to reward stakes. Vice versa, lower performing participants might expect occasional punishments and thus also attend to punishment stakes. There was no significant correlation between task performance and the number of first fixations on rewards vs. punishments, *r*(97) = −0.11, 95% CI [-0.30, 0.09], *p* = .298 and the association between task performance and the attentional effect on responses remained significant when controlling for the number of first fixations, *t*(96) = 4.97, *p* < .001. There was however though a significantly negative correlation between task performance and overall dwell time difference (dwell time on reward stakes minus dwell time on punishment stakes), *r*(97) = −0.27, 95% CI [-0.44, −0.08], *p* = .007: participants with higher performance showed a more variable (i.e., less biased towards reward stakes) gaze pattern and attended relatively more to punishments compared to participants with low performance. The association between task performance and the attentional effect on responses remained significant when controlling for the this overall dwell time difference, *t*(96) = 5.20, *p* < .001. In sum, we found no evidence for high performing participants exclusively focusing on reward stakes and low performing participants also attending to punishment stakes. If anything, we found the opposite pattern of high performing participants showing a more variable gaze pattern (also attending to punishment stakes), which chimes with the idea that these participants could rely their response on their (more adaptive/ flexible) gaze pattern.

Note that all these performance-dependent results are exploratory and should be interpreted with caution.

**Figure SI07.**
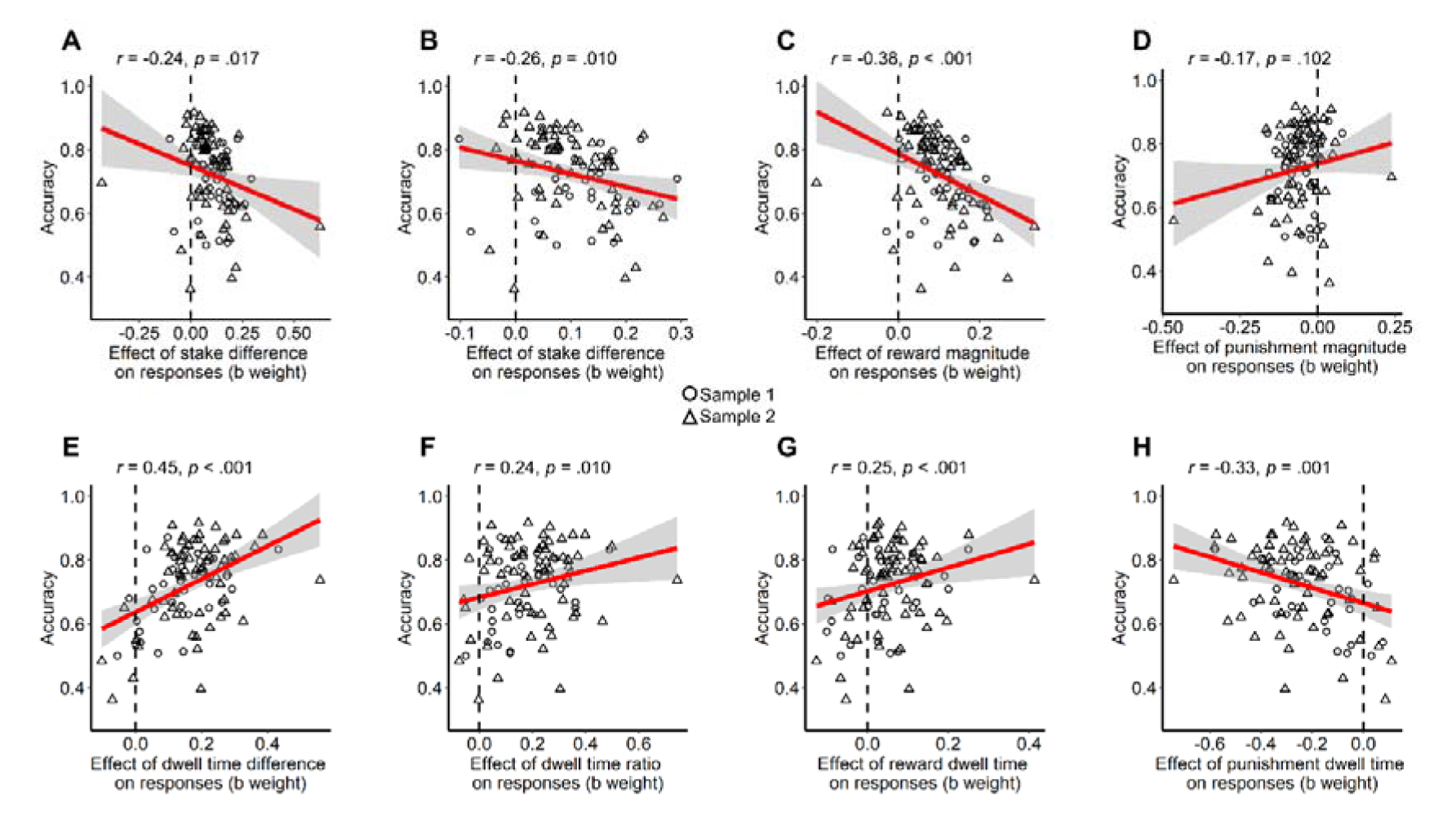
Association between interindividual variability of accuracy and in the effects of stake magnitudes and dwell times on responses. Participants’ mean accuracy correlated significantly negatively with their respective effect of stake differences on responses (**A**), also when two outliers removed (**B**), which was driven both by a negative correlation with the effect of the reward stake (**C**; note that these effects tend to be positive) as well as a positive correlation with the effect of the punishment stake (**D**; note that these effects tend to be negative, i.e., participants with stronger negative effects showed worse performance). These correlations suggest that participants with strong stake difference effects showed poor performance. The opposite pattern occurred for the effect of dwell time on responses: This effect correlated significantly positively with accuracy, both for the difference between reward and punishment dwell times (**E**) as well as the relative dwell time (ratio) on rewards (**F**). Again, this effect was driven by reward dwell times (**G**) rather than punishment dwell times (**H**). These correlations suggest that participant with strong attention effects showed high performance.

